# Nucleotide State Regulates Asymmetric Substrate-Binding Protein Docking in a Type II ABC Importer

**DOI:** 10.64898/2026.10.15.759412

**Authors:** Alexandra Karagiaridi, Ramon Mendoza Uriarte, Muyuan Chen, Benoît Roux, Heather W. Pinkett

## Abstract

Type II ABC importers rely on substrate-binding proteins (SBP) to capture nutrients and deliver them to membrane-embedded transporters, but how ATP binding and hydrolysis regulate their interactions remains poorly understood. To address these questions, we determined structures of the Type II small-substrate molybdate importer MolBC-A from *H. influenzae* in distinct nucleotide-bound states, complemented by molecular dynamics (MD) simulations and functional assays. In a nucleotide-free state, the MolA-MolBC interface is remodeled into a more open conformation, whereas ATP and AMP-PNP binding induce asymmetric conformational changes that preserve the MolA Lobe I interface and disrupt contacts with MolA Lobe II, revealing a previously unresolved partially-docked state. In the ADP-bound state, MolA exhibits the greatest conformational heterogeneity with reduced stable engagement that extends across both lobes. Together these findings define a nucleotide-dependent cycle of asymmetric MolA remodeling and reveal distinct contributions of the two SBP lobes in coupling transporter nucleotide state to SBP engagement.

## Introduction

ATP-binding cassette (ABC) transporters are a large family of membrane proteins that facilitate the unidirectional transport of molecules across the membrane via ATP-mediated conformational changes. ABC transporters function as either importers or exporters and participate in processes ranging from nutrient uptake to cellular defense and regulation^1–4^. Despite their functional diversity, canonical ABC transporters share a conserved core architecture comprised of two transmembrane domains (TMDs), which form the translocation pathway, and two nucleotide-binding domains (NBDs), which bind and hydrolyze ATP^5–7^. In ABC importers, architectural differences in the TMDs further subdivide them into Type I, Type II, and energy-coupling factor (ECF), which employ unique mechanisms of substrate recognition and translocation^5,8–13^. Type I and Type II importers, in particular, rely on an additional substrate-binding protein (SBP) to capture and deliver substrates to the transporter.

Substrate recognition by the SBP and the conformational changes accompanying binding provide an initial layer of transport selectivity in Type I and Type II importers. Despite low sequence conservation among SBPs with different substrate specificities, these proteins share a conserved topology of two lobes connected by a flexible hinge region. The binding pocket is located at the interface between the two lobes, and substrate-binding shifts the SBP between conformational states that influence its interaction with the transporter^14,15^. Canonically, closure of the SBP has been regarded as favorable for a productive association with the transporter. However, multiple competent SBP conformations have been observed, including ones that differ by only a few degrees of closure even when bound to similar substrates, suggesting the SBP-transporter docking mechanism is not dictated by a single SBP closed-state geometry. Coupled with the ability of some SBPs to bind non-cognate substrates and still interact with their cognate transporters, but not necessarily support transport, suggests that complex formation may provide an additional layer of transport selectivity^16^.

Transport therefore requires a substrate-loaded SBP conformation that is compatible with the docking state of the transporter. In Type I importers, this is initiated by substrate-binding leading to ligand-induced SBP closure that is distinct enough from the apo conformation and favors transporter association. Docking is coupled to ATP-dependent NBD closure, where ATP binding and hydrolysis then alternate the TMDs between outward– and inward-facing conformations to support substrate release^17,18^. Substrate-binding sites within the transporter can provide an additional layer of selectivity, distinguishing ligands recognized by the SBP from those that are efficiently transported^19,20^. Type II importers appear to use a distinct coupling mechanism. Their SBPs can associate tightly with their nucleotide-free transporters in absence of substrate, while nucleotide-binding and hydrolysis can destabilize SBP-transporter interactions to varying extents across Type II systems^21–25^. With Type II systems also lacking a highly selective substrate-binding cavity, specificity may reside primarily in the SBP or in additional, as yet unresolved features of the translocation pathway, such as steric constraints imposed by pore size. Because Type II importers transport substrates that vary widely in size and chemical properties, how SBP recognition is coupled to transporter docking and translocation remains incompletely understood.

To define how nucleotide state regulates SBP engagement during the transport cycle, we determined structures of the small-substrate molybdate MolBC-A importer from *H. influenzae* in distinct nucleotide states and in absence of bound substrate. MolBC consists of the MolB transmembrane (TMDs) and MolC nucleotide-binding domains (NBDs) and associates with the substrate-binding protein (SBP) MolA via an unknown mechanism. Our data support a nucleotide-dependent model in which ATP binding forms a previously unresolved partially docked state, with MolA anchored to the transporter through one lobe, while interactions with the second lobe are destabilized. In the ADP-bound state MolA shows the greatest conformational heterogeneity and reduced stable engagement with MolBC, with conformational changes involving both lobes. This structural framework provides new insights into how asymmetric SBP engagement is coupled to the catalytic cycle of Type II ABC importers.

## Results

### MolA docking locally remodels the periplasmic gate of MolBC

To determine how SBP docking influences the architecture of a small-substrate Type II importer, we determined the nucleotide-free cryo-EM structure of *H. influenzae* MolBC in complex with substrate-free MolA to 3.15Å resolution (PDB ID:38RC). MolA docks onto the periplasmic surface of MolBC and induces localized structural changes at the MolA-MolBC interface. Superposition of MolBC (PDB ID:38MO) with the nucleotide-free MolBC-A complex (RMSD=0.633Å) shows that MolA binding does not substantially alter the open conformation of MolC nucleotide binding domains (NBDs) **(Fig. 1A)**. MolA docking also does not change the overall inward-facing conformation of the translocation pathway formed at the interface of the MolB (TMDs) dimers. This differs from the SBP-bound *E. coli* vitamin B_12_ BtuCD-F importer but resembles the inward-facing conformation observed for the heme importer BhuUV-T from *B. cenocepacia*, both of which belong in the Type II ABC importer family^21,26^.

**Figure 1:**
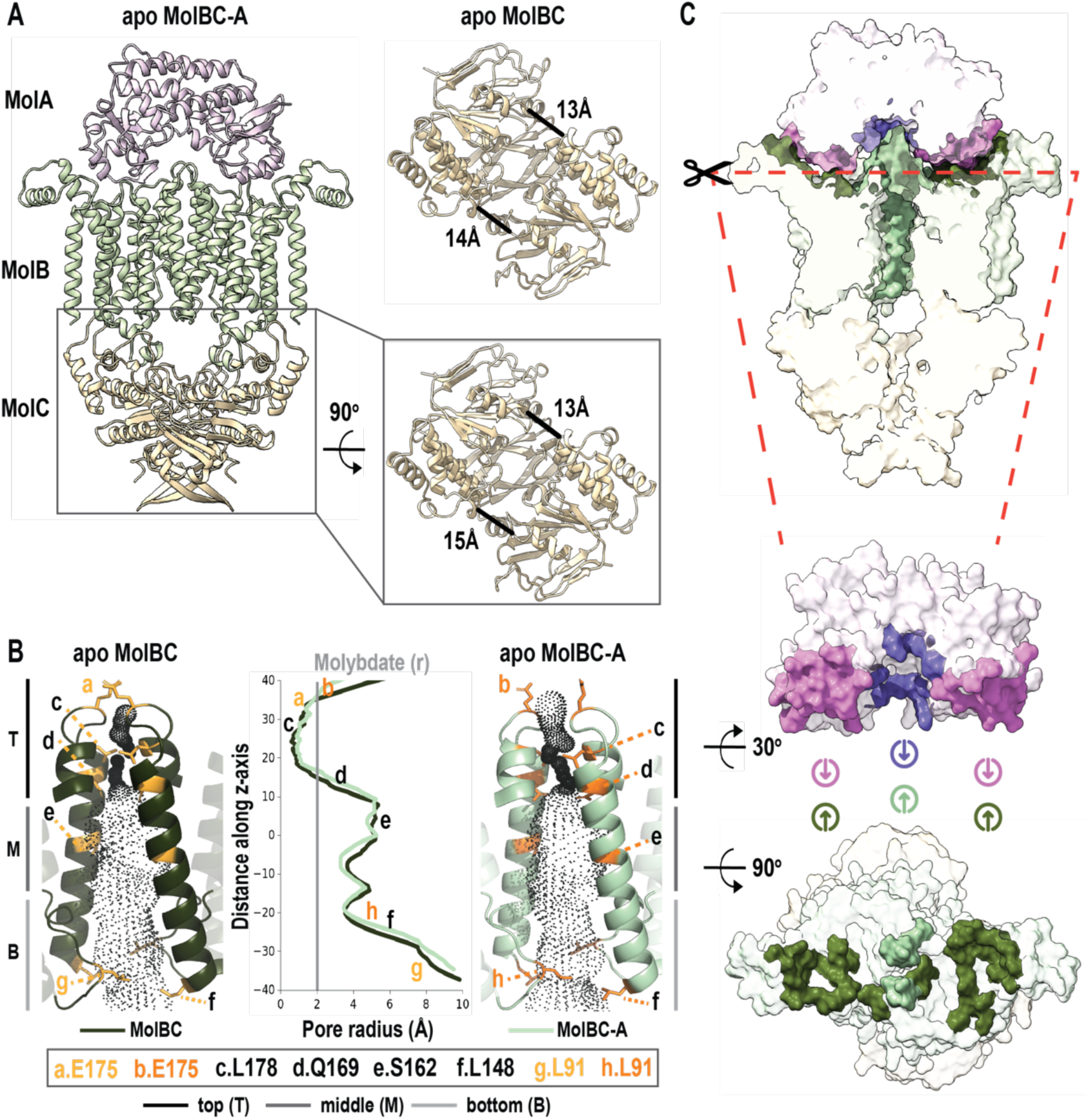
MolA docking locally remodels the periplasmic gate of MolBC. **A**, Structure of nucleotide-free (apo) MolBC-A transporter with a top view zoom into the dimerized MolC subunits (NBDs; wheat, bottom). Enlarged cytoplasmic views compare the MolC NBDs of MolBC-A and MolBC alone. Cα-Cα distances between S130 in the signature motif and C42 in the Walker A region of the opposing subunits are indicated for both NBDs. **B,** Comparison of the translocation pathway of nucleotide-free MolBC (apo; left; dark green) and nucleotide-free MolBC-A (apo; right; light green). TM5 and TM5α, which line the central pathway, are shown in cartoons, with the pore dimensions calculated using HOLE represented by black points. The central plot shows pore radius along the membrane-normal z-axis from the periplasmic (top) to the cytoplasmic side (bottom). E175 marks the periplasmic entrance, L178 the deeper TM5α constriction, Q169, and S162 the midpoint of the pathway, and L148 and L91 cytoplasmic gates I and II, respectively. **C,** Surface representation (top) of the MolBC-A docking interface. MolBC-A shown in white with interacting residues and translocation pathway highlighted by color to show asymmetric contacts formed by MolA Lobe I and Lobe II (pink, purple) with the two MolB subunits (dark and light green). The MolBC surface has been clipped to visualize the translocation pathway (light green). Below MolA and MolBC are shown in a 30° bottom view and 90° top view, respectively, to highlight the asymmetric contacts.

The central translocation pathway is lined primarily by TM5 and TM5α from each MolB subunit, with TM5/5α loops forming the periplasmic entrance. Access to the cytoplasm is facilitated by cytoplasmic gates I and II, formed by TM5 and TM2/3 loops, respectively, at the cytoplasmic side of the pathway. Closer inspection of the pathway shows that the effects of MolA docking are concentrated at the periplasmic entrance. MolA docking stabilizes the otherwise flexible TM5/5α loops in each MolB subunit, increasing the pore radius near E175 by ∼1-2Å, creating an opening sufficient to accommodate molybdate at the top of the periplasmic gate **(Fig. 1B: site b)**. Immediately below this opening, however, the constriction formed by the TM5α helices remains narrow with a minimum pore radius of ∼1Å (near L178), insufficient to accommodate molybdate **(Fig 1B: site c)**. Beyond this constriction, the pathway expands to approximately 3-6Å in radius, with both cytoplasmic gates remaining open **(Fig. 1B: sites d-f,h)**. Thus, MolA docking does not induce a global conformational transition in MolBC, but instead remodels and stabilizes the periplasmic entryway while maintaining the constriction within the TM5α region of the pathway. These localized changes were not resolved in previous EPR measurements, where the relevant region was sampled by a limited number of labeled sites^27^.

Outside of the translocation pathway, MolA binding is coupled with rotation of the short helices connecting TM1 and TM2 on the periplasmic side of each MolB subunit. These helices are poorly resolved in MolBC alone but become well-ordered in the MolBC-A complex, indicating that MolA stabilizes this region upon docking. MolA engages each MolB subunit through a network of predominantly charged interactions, with more extensive contacts formed by the N-terminal lobe (Lobe I) of MolA. These interactions span across the MolB periplasmic pocket, whereas interactions involving the C-terminal lobe (Lobe II) are confined to a smaller region of the opposite side of the pocket **(Fig. 1C)**. Notably, MolB residues R62 and R65 on one subunit form contacts with E99 on MolA Lobe I, while the same residues on the other subunit contact E249 on MolA Lobe II. Corresponding charged residues are conserved among Type II importers, and mutations at analogous positions have been linked to impaired transport or destabilization of the SBP-transporter complex^21,26,28^. Collectively, our MolBC-A structural observations suggest that the two MolA lobes make different contributions in stabilizing specific periplasmic elements of MolBC.

### MolBC binding selectively remodels MolA Lobe II and the substrate-binding pocket into a more open conformation

Complex formation also induces asymmetric changes in MolA that remodel its substrate-binding pocket. A comparison of MolBC-A with the structures of holo-MolA bound to molybdate (PDB ID:3PSH; RMSD=0.914Å) or tungstate (PDB ID:3PSA; RMSD=0.893Å) shows that Lobe I remains largely unchanged, while the lower region of Lobe II undergoes an outward displacement of approximately 7Å in the MolBC-A complex. Neighboring surface loops and β-strands move in the same direction, producing a localized opening of the substrate-binding pocket **(Fig. 2A)**. Closer inspection of MolA within the substrate-free MolBC-A complex (more open conformation) and isolated, substrate-bound MolA (closed conformation) shows that complex formation also disrupts a subset of Lobe II substrate-binding contacts **(Fig. 2A)**. In isolated, substrate-bound MolA, K221 and R264 contribute to coordination of the bound ligand, whereas in the substrate-free MolBC-A complex these residues instead interact with periplasmic elements of MolB, including the TM5/5α region containing E175 at the substrate entry site **(Fig. 2B)**. Additional MolBC-A contacts include Y301 in Lobe IΙ and Y114 in Lobe Ι, both of which interact with the MolB periplasmic loops, but do not undergo comparable rearrangement in the binding pocket. As a result, MolBC binding selectively remodels Lobe II and positions the periplasmic loops of MolB within the MolA binding pocket, disrupting a subset of substrate-coordinating interactions while leaving Lobe I largely unchanged.

**Figure 2:**
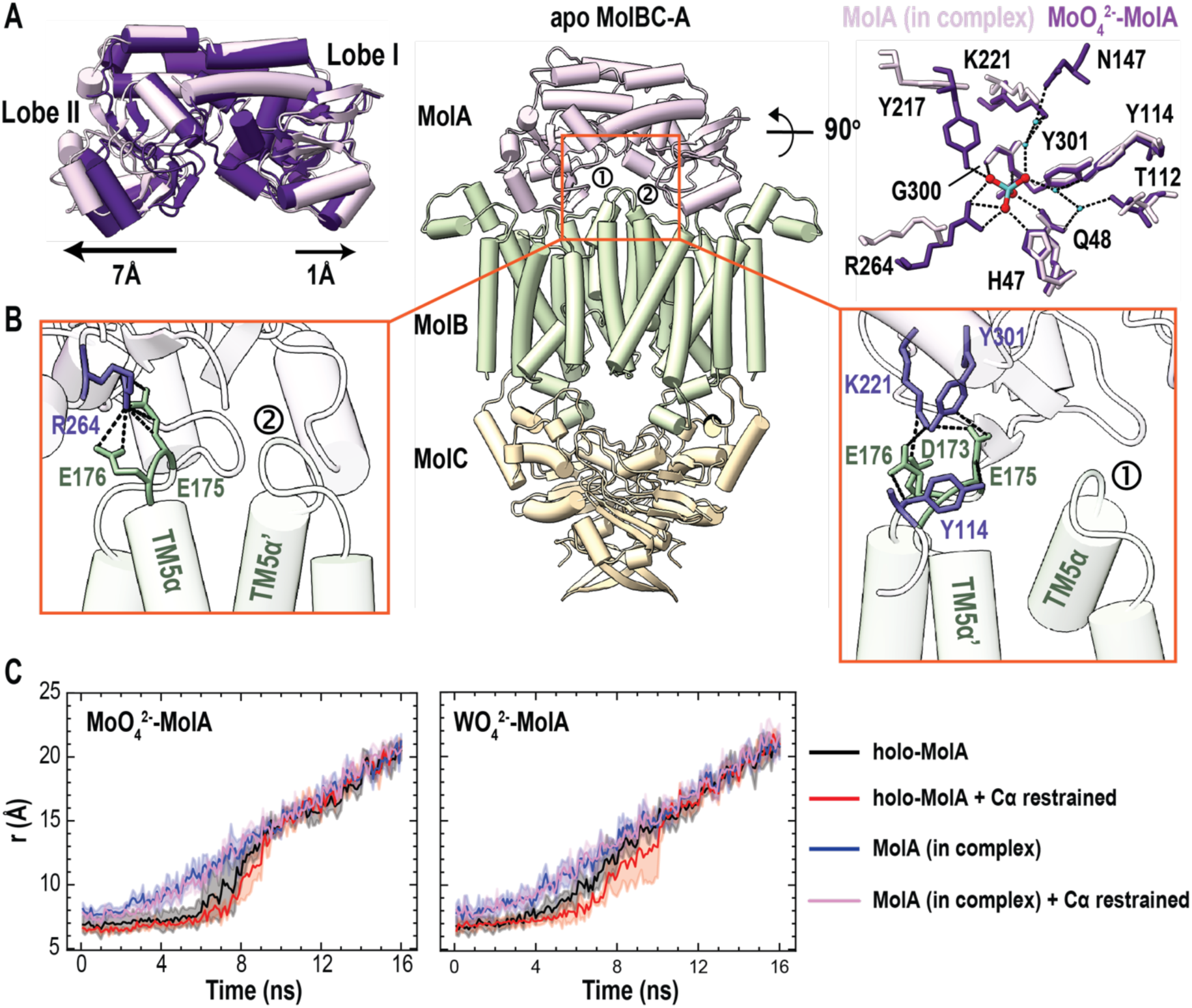
MolBC binding remodels MolA Lobe II and the substrate-binding pocket. **A**, Structural alignment of molybdate-bound isolated MolA (PDB ID: 3PSH; purple) and substrate-free MolA from nucleotide-free MolBC-A (apo; light pink), showing the transporter-induced outward displacement of Lobe II, while Lobe I remains largely unchanged. On the far right, a bottom view of the MolA substrate-binding pocket is shown with residues involves in molybdate binding as sticks. The MolBC complex is omitted for clarity. **B,** Interactions between MolA binding pocket (purple) and the periplasmic loops of the two MolB subunit (green) that form the entrance to the translocation pathway. Portions of MolA and MolB are omitted for clarity, and interacting residues shown as sticks. **C,** Distance between the center of masses of MolA backbone and substrate; the MolA Cα atoms observed in MolBC-A or the closed, substrate bound (holo-MolA) structure were restrained in pink and red and not restrained in blue and black, respectively. The left and right plots show unbinding of molybdate or tungstate, respectively, over the 15ns simulation. Each trace is the average of four independent simulations (n=4), with standard deviation represented by the gray shaded area.

To assess whether transporter-induced remodeling of MolA affects substrate retention, we compared MolA in a closed (substrate-bound) and the more open conformation observed in the MolBC-A complex using molecular dynamics (MD) simulations. MolBC was omitted from these simulations so that the effects of the two MolA conformations could be compared directly. For the more open conformation, molybdate or tungstate was positioned in the binding pocket by superposition with the corresponding closed, substrate-bound MolA structure. In the trajectories, both molybdate and tungstate were displaced from the more open conformation earlier along the pulling trajectory (t=4ns) than from the closed conformation (t=6ns) **(Fig. 2C)**. Consistent with this difference, the force required to displace either substrate from its binding pocket was approximately 4-fold lower from the more open conformation **(Supplementary Fig. 1)**. Thus, under equivalent pulling conditions, the MolBC-A derived MolA conformation offers less resistance to substrate displacement than closed MolA. Coupled with the complex-induced loss of Lobe II substrate-coordinating contacts and their replacement by interactions with the MolB periplasmic gate loops, these results support a model in which transporter docking remodels the MolA binding pocket into a conformation more permissive for substrate release.

### ATP binding allosterically narrows cytoplasmic gate II

To determine whether ATP binding drives structural rearrangements beyond those induced by MolA docking, we solved the cryo-EM structure of ATP-bound MolBC-A at 2.74Å resolution (PDB ID:38MT). To stabilize the ATP-bound state, the catalytic glutamate in MolC (E154) was mutated to glutamine, an established approach for reducing ATP hydrolysis in ABC transporters^17,29,30^. Well-resolved density allowed ATP/Mg^2+^ to be modeled at both NBD interfaces, with each nucleotide coordinated by the signature motif (LSGGQ) of one MolC subunit and the Walker A/B motifs and A-, H-, and Q-loops of the opposing subunit **(Fig. 3A)** consistent with canonical ABC transporter nucleotide-binding sites^2,17,31,32^.

**Figure 3:**
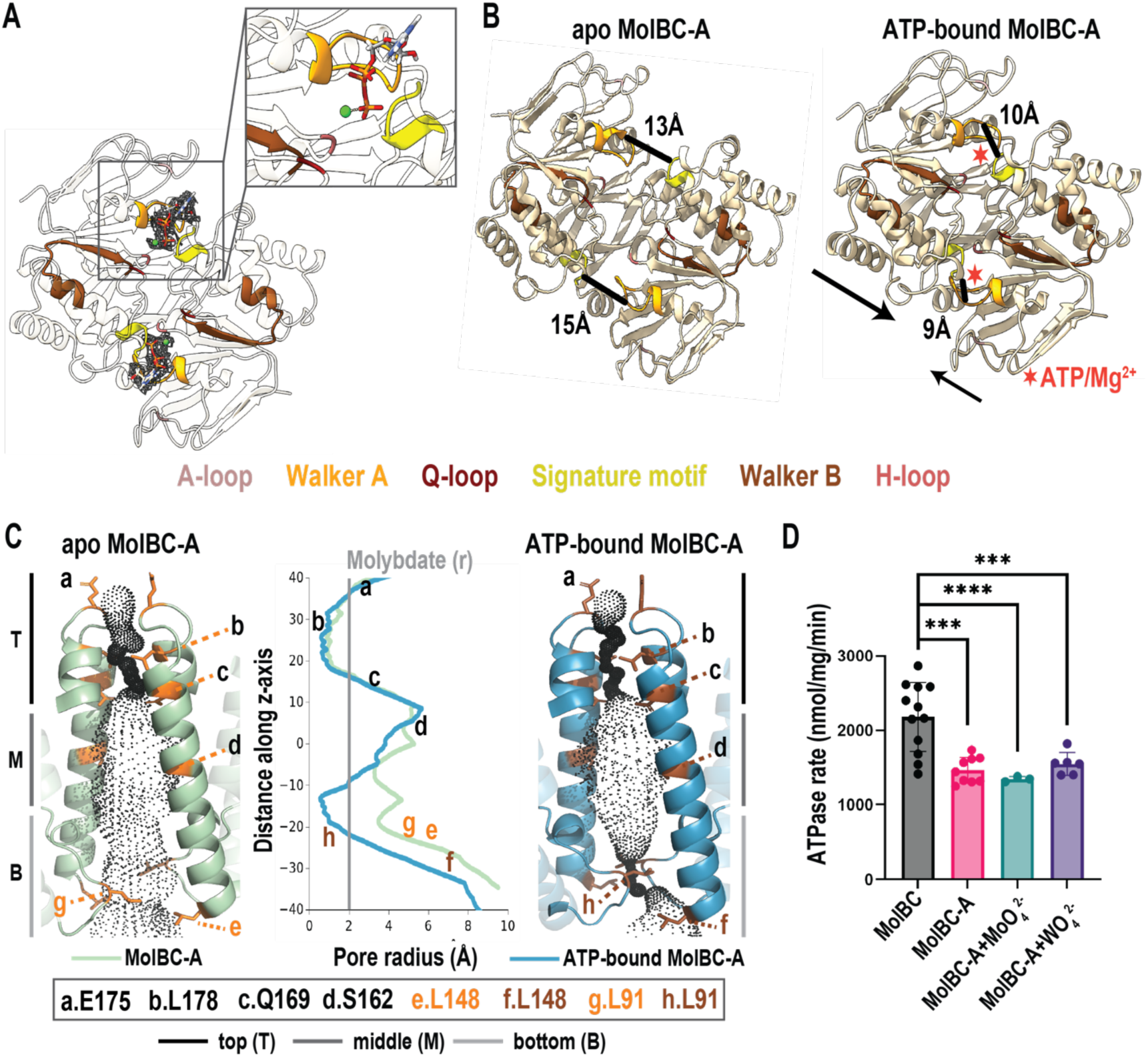
ATP binding initiates asymmetric transporter closure and narrows cytoplasmic gate II. **A**, Enlarged top view of the ATP binding sites in ATP-bound MolBC-A. ATP shown as sticks and magnesium ions as green spheres, with corresponding density shown around each ligand. Conserved nucleotide-binding motifs are colored according to the key. **B,** Cytoplasmic (top) views of the dimerized MolC NBDs (wheat) in nucleotide-free MolBC-A (apo; left) or ATP-bound (right) MolBC-A, showing ATP-induced NBD closure. Cα-Cα distances between S130 in the signature motif of one MolC subunit and C42 in the Walker A motif of the opposing subunit are indicated for both NBDs. Arrows indicate the unequal movements of the two MolC subunits upon ATP binding. Red asterisks mark the positions of ATP/Mg^2+^, which are omitted for clarity. **C,** Comparison of the translocation pathway of nucleotide-free MolBC-A (apo; left; light green) and ATP-bound MolBC-A (right; blue). TM5 and TM5α, which line the central pathway, are shown in cartoons, with the pore dimensions calculated using HOLE represented by black points. The central plot shows pore radius along the membrane-normal z-axis from the periplasmic (top) to the cytoplasmic side (bottom), as in Fig. 1B. **D,** ATPase activity of MolBC in the absence or presence of MolA, molybdate or tungstate. Bars represent mean ATP hydrolysis rates from at least three technical replicates (MolBC n=12, MolBC-A n=9, MolBC-A+MoO_4_^2-^ n=3, MolBC-A+WO_4_^2-^ n=6); error bars indicate standard deviation. Statistical significance values determined by Welch’s t-test are indicated above bars.

ATP binding induces asymmetric closure of the MolC NBDs that is coupled primarily to the narrowing of cytoplasmic gate II, while the remainder of the MolB translocation pathway remains similar to that of the nucleotide-free MolBC-A complex. Alignment of the nucleotide-free and ATP-bound MolBC-A structures (RMSD=0.949Å) reveals an approximately 8Å NBD closure, with one MolC subunit undergoing a larger displacement than the other **(Fig. 3B)**. Unlike the more extensive TMD rearrangements associated with nucleotide binding in Type I importers, these asymmetric movements in ATP-bound MolBC-A are concentrated at the cytoplasmic ends of the transmembrane helices. Consequently, cytoplasmic gate II narrows, while leaving cytoplasmic gate I and the periplasmic gate largely unchanged. HOLE analysis quantitatively supports the localized effect of ATP binding on cytoplasmic gate II. Relative to the nucleotide-free MolBC-A, ATP binding narrows the pore near L91 by approximately 4Å, converting this region from the widest portion of the cytoplasmic gate in the nucleotide-free structure, to the narrowest when ATP-bound **(Fig. 3C: site h)**. Conversely, the pore radii at cytoplasmic gate I (L148) and periplasmic gate radii (E175, L178, Q169) remain largely unchanged **(Fig. 3C: sites a-d, f)**. These structural changes are consistent with our previous EPR measurements showing nucleotide-dependent closure of gate II^27^. However, TM5α region remains constricted in the ATP-bound structure with a minimum pore radius of ∼0.8Å near L178, too narrow to accommodate molybdate. Thus, the ATP-bound cryo-EM structure shows selective narrowing of the cytoplasmic side of the pathway but does not capture the increased opening or flexibility of the periplasmic gate, as inferred from EPR^27^.

Consistent with the persistent TM5α constriction, no tungstate density was detected despite its inclusion during structure preparation. Because substrate was not resolved structurally, we next examined whether molybdate or tungstate altered MolBC-A ATPase activity. MolA decreased ATP hydrolysis by approximately 2-fold relative to MolBC, whereas addition of molybdate or tungstate did not produce a further measurable decrease in activity. Increasing substrate concentration likewise did not further reduce ATPase activity, indicating that under these conditions it is difficult to distinguish substrate specific inhibition from effects beyond the inhibition associated with MolA docking **(Fig. 3D)**.

### Nucleotide state regulates MolA’s conformational heterogeneity

Previous studies of Type II importers have shown that nucleotide state can destabilize the SBP-transporter interactions, although the structural basis for this effect is unclear^22,23,25^. In the ATP-bound MolBC-A dataset, 98% of particles retained after 2D classification contained MolA density, although MolA Lobe II density was less well resolved compared to Lobe I. As a result, the final reconstructed atomic model captures a predominantly SBP-bound but partially docked state, where ATP selectively disrupts the MolA Lobe II interface while maintaining interactions with Lobe I. Compared to nucleotide-free MolBC-A, ATP-dependent asymmetric narrowing of the TM6α/β coupling helices is accompanied by poorer resolution of the periplasmic regions of TM1 and TM2 (residues 5-62), consistent with increased conformational heterogeneity. The short helices connecting TM1 and TM2, which contribute to the MolA docking interface, are consequently also unresolved. Additional rearrangements within the TM9-TM10 and TM5α-TM6 periplasmic loops disrupt contacts made by conserved arginine residues involved in MolBC-A complex assembly, resulting in further loss of interactions with MolA **(Fig. 4A)**. Together, these changes selectively destabilize the Lobe II side of the SBP-transporter interface, while maintaining Lobe I association, providing a structural basis for the asymmetric, partially docked MolA state.

**Figure 4:**
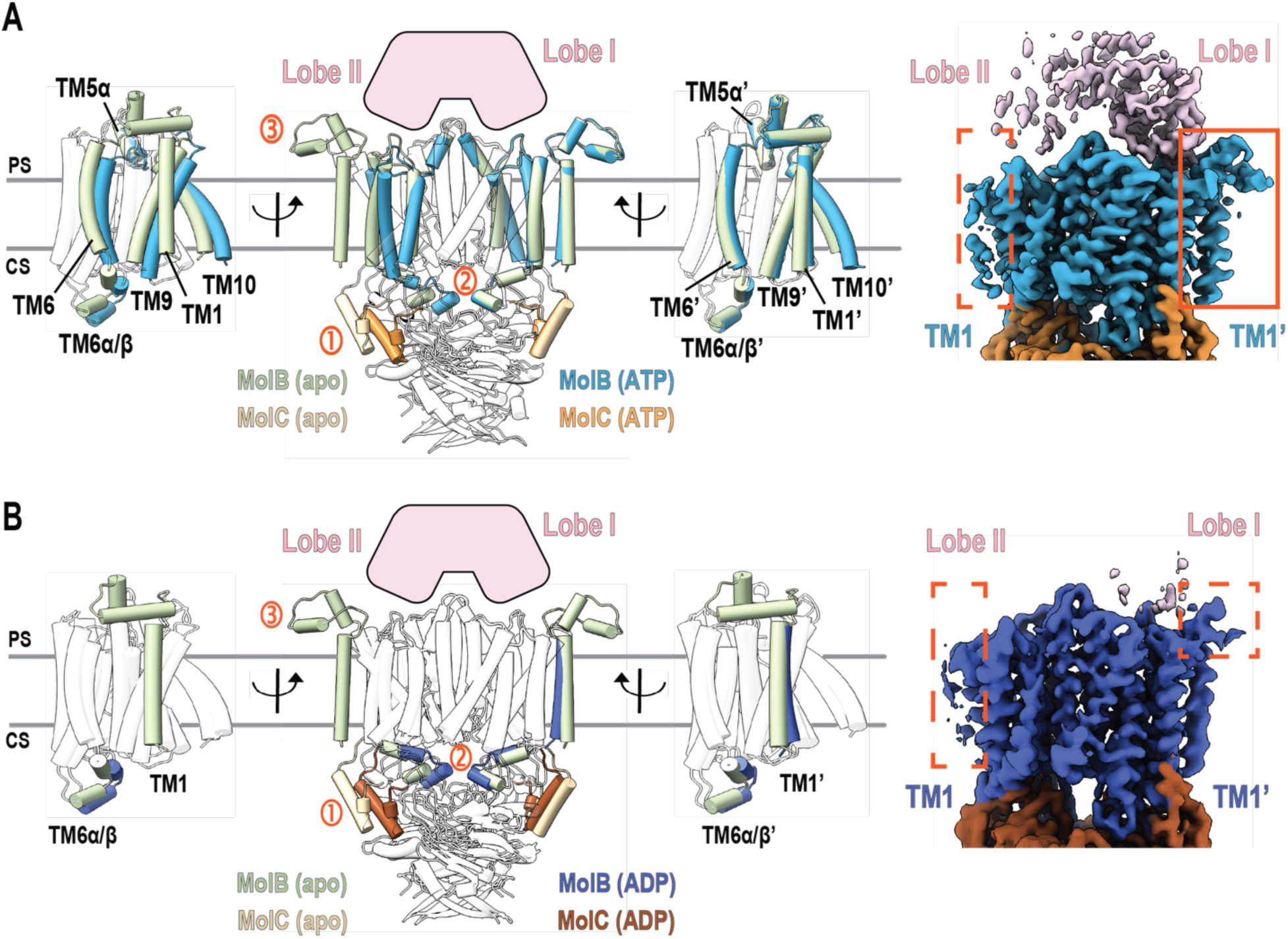
ATP binding asymmetrically destabilizes MolA docking, whereas the ADP-bound state favors disengagement. **A**, Front (middle) and side (left and right) views of nucleotide-free MolBC-A (apo; light green) with ATP-bound MolBC-A (blue), highlighting the asymmetric ATP-dependent rearrangement of relevant TM helices that disrupts contacts with MolA Lobe II. While 98% of 2D classified particles retain MolA density (pink cartoon), it is more poorly resolved in Lobe II vs. Lobe I precluding reconstruction of that side of MolA. Numbered labels in the front (middle) transporter view indicate: 1) asymmetric closure of the NBDs; 2) asymmetric decrease in distance between the two coupling-helices, TM6α/β region; 3) loss of contacts with MolA Lobe II. ATP and Mg^2+^ from each MolC subunit are hidden for clarity. Enlarged cryo-EM map view on the right shows loss of the MolA-MolBC interface on one side of the complex (dashed orange box), while contacts are retained on the opposite side (solid orange box). **B,** Same transporter views as in **A**, comparing nucleotide-free MolBC-A (apo; light green) with ADP-bound MolBC-A (dark blue). In the ADP-bound structure, the TM1-2 connecting helices are unresolved, and 2D particle classification shows that 56% of the particles lack a stably associated MolA. Numbered labels in the front (middle) transporter view correspond to the same structural changes as in **A**. ADP from each MolC subunit is hidden for clarity. Enlarged cryo-EM map view on the right shows enhanced disengagement of MolA from MolBC on both sides of the interface (dashed orange boxes).

To determine whether nucleotide binding or hydrolysis is associated with additional changes in MolA conformational heterogeneity, we determined the structures of MolBC-A bound to the non-hydrolysable ATP analogue, AMP-PNP (PDB ID:38RB) at 2.62Å resolution, and ADP (PDB ID:38RD) at 2.91Å resolution. Although all three nucleotide-bound structures show similar degrees of NBD closure **(Supplementary Fig. 2A, 2B)**, differences in their TMD conformations are associated with distinct degrees of MolA engagement. Despite these differences, HOLE analysis shows that the overall translocation pathway remains similar across all nucleotide-bound states. The periplasmic gate remains comparably open to the same extent as ATP in pre-(AMP-PNP) and post-(ADP) hydrolysis states **(Supplementary Fig. 3, sites a-d)**. Cytoplasmic gate I also is open to the same degree as ATP in the AMP-PNP-bound structure but narrows by 2Å in radius in the ADP-bound structure, without, however, altering the overall open conformation of that side of the pathway **(Supplementary Fig. 3, sites e and f)**. Finally, cytoplasmic gate II narrows in both AMP-PNP– and ADP-bound structures to the same degree as in the ATP-bound structure **(Supplementary Fig. 3, site g)**. In contrast, the degree of asymmetric narrowing of the TM6α/β coupling helices differs between nucleotide states reflecting differences at the MolBC-A interface. In the AMP-PNP complex, TM1 is better resolved than in the ATP-bound structure, whereas the short helices between TM1 and TM2 and periplasmic portion of TM2 remain poorly resolved as in the ATP-bound structure. Based on this TMD orientation, MolA still remains partially docked on MolBC via its Lobe I, with 99% of particles retained after 2D classification containing MolA density. In the ADP-bound dataset, however, MolA exhibits substantially greater conformational heterogeneity, with 44% of classified particles containing MolA density and 56% lacking stably resolved MolA. In the final atomic model, the TM6α/β coupling helices narrow from both TMD sides resulting in poorer resolution of the periplasmic elements in both MolB subunits. This is further accompanied by substantially greater bilobed MolA heterogeneity preventing reconstruction of a stably docked MolA state **(Fig. 4B)**.

To characterize the conformational heterogeneity associated with MolA, we performed focused alignment using the Gaussian mixture modeling GMM-based method in EMAN2. Refined particle orientations improved local resolution in the MolA Lobe II region, and MolA motion trajectories were generated by calculating the per-particle relative orientation between MolA and MolBC, allowing the visualization of the ensemble of MolA states. In the ATP-bound ensemble, most of the conformational heterogeneity is concentrated in MolA Lobe II, which undergoes a largely rigid-body displacement relative to Lobe I between a more open and closed conformation; Lobe I and the inter-lobe hinge show substantially smaller movements **(Fig. 5, Supplementary Movie 1)**. AMP-PNP produces a similar conformational ensemble, although the range of Lobe II orientations is more restricted than in the ATP-bound state **(Fig. 5, Supplementary Movie 2)**. In contrast, conformational heterogeneity in the ADP-bound ensemble extends to both MolA lobes and includes substantially greater conformational range **(Fig. 5, Supplementary Movie 3)**. Together, these analyses show that nucleotide state regulates the conformational range of each MolA lobe, with ATP and AMP-PNP primarily affecting Lobe II, and ADP exhibiting broader conformational heterogeneity involving both lobes.

**Figure 5:**
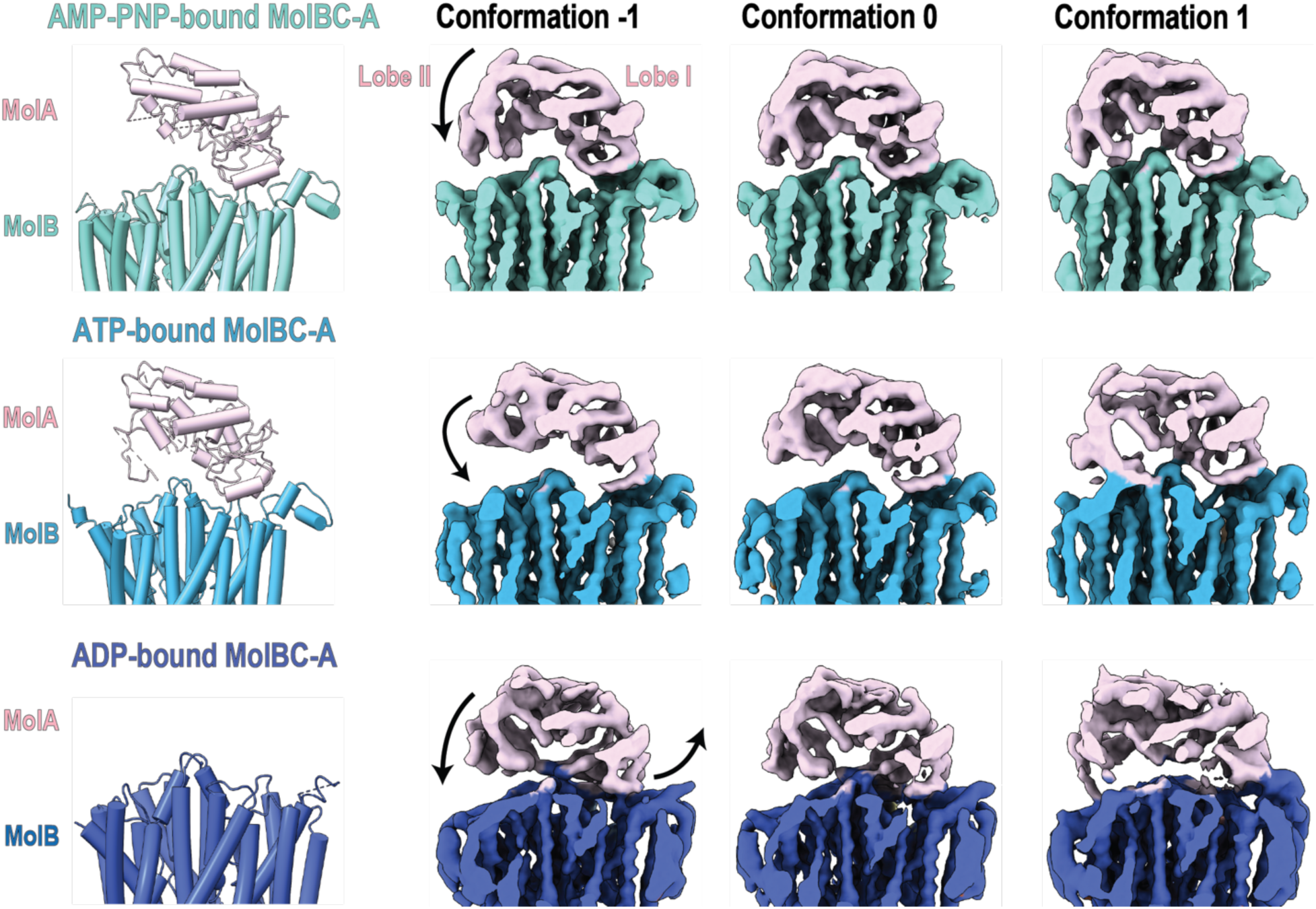
Nucleotide state reshapes the conformational ensemble of MolA. Representative composite maps illustrating MolA conformational heterogeneity in AMP-PNP-bound (top), ATP-bound (middle) or ADP-bound (bottom) MolBC-A. Maps were generated following focused refinement of MolA relative to MolBC using GMM-based analysis in EMAN2. Representative conformations illustrate the range of MolA orientations sampled within each nucleotide state. Arrows in Conformation –1 of each map depict the motion of the lobes relative to the rest of the transporter. The apparent amount of MolA density in these representative map views reflects the 2D classified percent of MolA-containing particles in each dataset: 99% in AMP-PNP-bound MolBC-A, 98% in ATP-bound MolBC-A, and 44% in ADP-bound MolBC-A.

To determine whether the nucleotide-dependent structural changes in MolA engagement were reflected in bulk binding kinetics, we measured the MolA-MolBC interactions by biolayer interferometry (BLI) in the presence and absence of nucleotide and substrate. In the absence of nucleotide, MolA bound MolBC with high affinity [KD=(1.28±0.01)’10^-9^M] characterized by a fast association rate [k_a_=4.43’10^5^ ± 7.14’10^2^ M^-1^s^-1^] and relatively slow dissociation [k_dis_=(0.6±0.005)’10^-3^ s^-1^]. Addition of ATP/Mg^2+^ did not measurably alter this interaction [KD=(1.26±0.02)’10^-9^M], consistent with our ATP-bound dataset, where majority of the particles reflect an SBP-bound state (2D classification: 98%). Mimicking of the post-hydrolysis nucleotide state with addition of ADP/Mg^2+^ also did not measurably alter the apparent affinity of MolA for MolBC [KD=(1.15±0.01)’10^-9^M], despite the substantially greater conformational heterogeneity observed in our ADP-bound cryo-EM dataset **(Table 1)**. Addition of molybdate or tungstate during the dissociation phase, in presence or absence of ATP/Mg^2+^, likewise did not accelerate MolA dissociation of the pre-assembled MolBC-A complex **(Table 1)**. These BLI results indicate that once the high affinity MolBC-A complex is assembled, it remains kinetically stable under the conditions tested. The absence of a measurable substrate-dependent effect is consistent with the nucleotide-free MolBC-A structure, in which periplasmic elements of MolB extend into and partially occlude the MolA substrate-binding pocket, potentially limiting substrate access to the assembled complex.

**Table 1:** Binding kinetic constants of MolA-MolBC interactions measured by BLI Octet.

| **Additives** | **$k_a$ ($M^{-1}s^{-1}$)** | **$k_{dis}$ ($s^{-1}$)** | **KD (M)** |
| --- | --- | --- | --- |
| None | $4.43 \times 10^5 \pm 7.14 \times 10^2$ | $(0.6 \pm 0.005) \times 10^{-3}$ | $(1.28 \pm 0.01) \times 10^{-9}$ |
| + ATP/Mg<sup>2+</sup> | $3.67 \times 10^5 \pm 8.66 \times 10^2$ | $(0.5 \pm 0.007) \times 10^{-3}$ | $(1.26 \pm 0.02) \times 10^{-9}$ |
| + ADP/Mg<sup>2+</sup> | $3.82 \times 10^5 \pm 6.94 \times 10^2$ | $(0.4 \pm 0.12) \times 10^{-3}$ | $(1.15 \pm 0.01) \times 10^{-9}$ |
| + MoO<sub>4</sub><sup>2-</sup> | | $(1.05 \pm 0.19) \times 10^{-3}$ | |
| + WO<sub>4</sub><sup>2-</sup> | | $(1.82 \pm 0.19) \times 10^{-3}$ | |
| + MoO<sub>4</sub><sup>2-</sup> + ATP/Mg<sup>2+</sup> | | $(1.17 \pm 0.46) \times 10^{-3}$ | |
| + WO<sub>4</sub><sup>2-</sup> + ATP/Mg<sup>2+</sup> | | $(0.61 \pm 0.29) \times 10^{-3}$ | |

## Discussion

Here, we present the first structural series of MolBC-A that captures distinct nucleotide-dependent states of the small-substrate Type II importer from *H. influenzae*. Together with GMM-based conformational analysis, these structures reveal nucleotide-dependent remodeling of MolA engagement with the transporter, that includes a partially docked SBP state, previously inferred from biochemical studies, but not structurally resolved, in which ATP binding preserves interactions with MolA Lobe I while disrupting contacts with MolA Lobe II^24,32^. Together, these findings support a model in which nucleotide-dependent conformational changes in MolBC allosterically remodel the SBP-transporter interface and regulate the extent of MolA engagement during the transport cycle **(Fig. 6)**.

**Figure 6:**
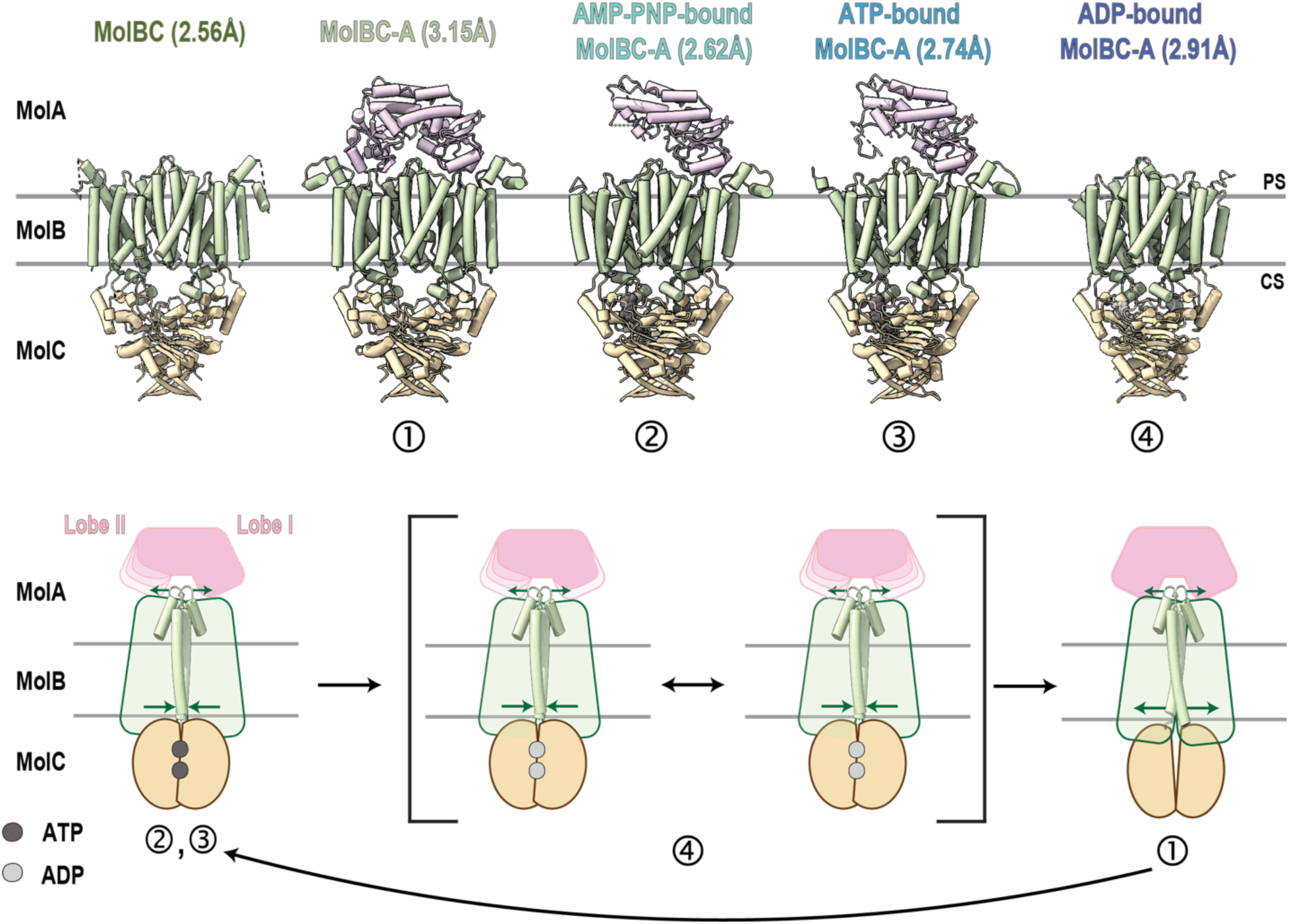
Proposed model of nucleotide-dependent MolA docking during the MolBC-A catalytic cycle. Top, Summary of the cryo-EM determined structures of *H. influenzae* MolBC-A captured in distinct nucleotide states, with corresponding resolutions indicated above each structure. PS and CS denote the periplasmic and cytoplasmic sides, respectively. **Bottom,** Model for nucleotide-dependent remodeling of the MolBC-A interface. ATP binding induces asymmetric closure of MolC NBDs and selectively disrupts the contacts between MolB and MolA Lobe II, while maintaining the Lobe I interface, producing a partially docked state (2,3). The ADP-bound state exhibits higher conformational heterogeneity affecting one or both lobes of MolA (4). Only in the nucleotide-free state, MolA can fully engage with MolBC through contacts involving both MolA lobes (1). Subsequent ATP binding re-establishes the asymmetric, partially docked MolA state (2,3). Green arrows indicate relative changes in the periplasmic and cytoplasmic gates observed across the nucleotide– and MolA-bound structural states.

In absence of nucleotide, MolA engages MolBC via interactions involving both lobes and the periplasmic surface of the transporter **(Fig. 6, atomic model 1)**. BLI confirms that this interaction is of high affinity **(Table 1)**, while the structure reveals asymmetric remodeling concentrated in MolA Lobe II that results in a more open conformation. Residues that coordinate substrate binding in isolated, substrate-bound MolA instead contact periplasmic elements of MolB in the complex, resulting in an extension of the periplasmic loops into the MolA binding pocket. MD simulations further suggest that substrates are displaced more readily from the more open, MolBC-A derived MolA conformation than the isolated, closed MolA, consistent with Lobe II remodeling weakening the interactions that retain substrate in the binding pocket. The insertion of periplasmic portions of MolB in the MolA binding pocket is consistent with the SBP docking mechanism of other Type II importers, suggesting a shared feature in SBP-transporter association^21,26^. In MolBC-A, the MolA-MolBC association also expands the periplasmic entrance near E175 on each MolB subunit sufficiently to accommodate a small substrate, like molybdate, although the deeper TM5α constriction remains too narrow. Collectively, our nucleotide-free MolBC-A structural observations and MD simulations suggest that asymmetric MolA docking remodels both the SBP-binding pocket and periplasmic entrance into a state that may facilitate substrate release and handoff.

GMM-based MolA focused refinement of the ATP-, AMP-PNP-, and ATP-bound states reveals conformational heterogeneity beyond that captured by the atomic models. The majority of the ATP– and AMP-PNP-bound particles retain MolA association with the transporter predominantly through Lobe I, while Lobe II samples a broader range of orientations **(Fig. 6, atomic models 2, 3)**. In contrast, the ADP-bound dataset samples a substantially broader range of MolA transitions involving one or both lobes, with 56% of 2D classified particles lacking a stably resolved MolA, density, precluding reconstruction of a stably docked MolA atomic model **(Fig. 6, atomic model 4)**. Conformational heterogeneity analysis further indicates that MolA Lobe I remains comparatively stable in the ATP– and AMP-PNP-bound states, whereas in the ADP-bound state both lobes undergo larger displacements and sample a broader conformational range. Together these observations suggest distinct contributions of the two MolA lobes to complex stability: one lobe providing a more stable docking interface while the other lobe is more responsive to substrate binding and nucleotide-dependent conformational changes in MolBC. Despite this structural remodeling, BLI measurements in the presence of ATP/Mg^2+^ or ADP/Mg^2+^ show no measurable reduction in the apparent affinity of the MolA-MolBC interactions relative to nucleotide-free conditions **(Table 1)**. The apparent differences likely reflect the distinct properties measured by each approach: cryo-EM and focused refinement resolve heterogeneity of MolA docking, whereas BLI reports ensemble-averaged association and cannot distinguish among differently engaged bound states. Stable MolA-MolBC association in presence of ADP/Mg^2+^ agrees with previously published SPR and pull-down experiments of other Type II importers^21–25^. In the same studies, addition of ATP/Mg^2+^ greatly destabilized, but did not completely abolish SBP-transporter association, whereas ATP/Mg^2+^ did not measurably alter the apparent affinity of MolA for MolBC under our BLI conditions.

Considering that intracellular ATP concentrations are typically high, the partially docked, single-lobe anchored state observed in ATP-or AMP-PNP-bound MolBC-A may be physiologically relevant. In Gram-positive bacteria, SBPs are often membrane anchored or fused to the transporter, limiting their diffusion away from the transporter, whereas SBPs in Gram-negative organisms are typically soluble periplasmic proteins^14,33^. Although MolA is not covalently tethered to MolBC in our structures, retention of one MolA lobe during the ATP-bound state could similarly maintain the SBP near MolBC during portions of the transport cycle. Under conditions where molybdate availability is limiting, such proximity could reduce reliance on repeated diffusion-limited encounters between MolA and the transporter^10,34^. Consistent with a potential compensatory role, MolBC-A is transcriptionally upregulated when the primary Type I transport system is inhibited in *H. influenzae*^35–37^. Partial retention of MolA with MolBC could therefore facilitate SBP re-engagement during successive transport cycles.

Finally, neither MolA nor ATP produce the canonical outward-facing conformation observed in Type I importers^17,34,38,39^. Instead, MolA docking locally expands the periplasmic entrance, whereas ATP binding primarily narrows cytoplasmic gate II. Despite these changes, the deeper TM5α constriction remains too narrow for molybdate passage in all nucleotide-bound states examined. Although none of our structures capture a fully open TM5α constriction, previous EPR measurements indicate increased nucleotide-dependent dynamics in this region. Because EPR reports conformational sampling, whereas cryo-EM preferentially resolves more highly populated structural states, these observations together suggest that an additional ∼1Å increase in pore radius at the TM5a constriction could occur transiently. Such a dynamic intermediate could permit substrate passage without requiring a long-lived globally outward-facing conformation. A comparable TM5α region forms the translocation bottleneck in Type II importers for heme and vitamin B_12_, although passage of these substantially larger substrates would likely require more extensive conformational rearrangements.

Together, these findings define asymmetric SBP docking engagement as a central feature of the MolBC-A catalytic cycle and reveal how nucleotide-dependent remodeling of the transporter reshapes SBP docking across distinct catalytic states. Determining how substrate occupancy alters this asymmetric interface and couples MolA engagement to transient opening of the TM5α gate will be important for understanding how substrate recognition is integrated with ATP-driven transport in Type II ABC importers.

## Methods

### Expression and purification of MolA and MolBC

The MolA binding protein was expressed and purified as described previously with the following modifications^35^. MolA (C-terminal 6xHistidine tag) was expressed in *E. coli* BL21(DE3) cells (EMD Biosciences) at 37°C in LB and harvested after a 2hr induction with 1mM IPTG. 2.5-5g of cells were lysed in osmotic shock solution (10mM Tris-HCl pH=7.5, 1mM EDTA pH=8.0 and 40% sucrose) with 1mM PMSF for 1hr at 4°C. Buffer components (25mM Tris-HCl pH=7.5, 2mM MgCl_2_, 250mM NaCl, 17mM Imidazole pH=8.0) were then added and the centrifuged supernatant was batched with Ni-NTA resin (Thermo Scientific) for 30min at 4°C. Following 10 column volume (CV) washes with wash buffer (25mM Tris-HCl pH=7.5, 250mM NaCl, 17mM Imidazole pH=8.0), bound MolA was eluted in 10CVs elution buffer (25mM Tris-HCl pH=7.5, 250mM NaCl, 150mM Imidazole pH=8.0) and dialyzed in 25mM Tris-HCl pH=7.5 and 250mM NaCl. For BLI, the dialysis buffer also included TEV protease at a 20:1 protein:TEV ratio, and cleaved MolA was purified via reverse affinity purification. Purified MolA was concentrated using a 30kDa MWCO concentrator (Millipore) and stored at –80°C for other experiments. For structural and biophysical studies with substrate, MolA was instead expressed in M9 minimal media and purified the same way.

The homodimeric membrane complex MolBC or the catalytically inactive mutant MolBC_E154Q_ were expressed and purified as before with the following modifications^40^. MolBC (N-terminal 6xHistidine tag) was expressed in E. *coli* BL21(DE3)plysS cells (EMD Biosciences) at 37°C in LB and harvested after a 3hr induction with 0.4mM IPTG. 5g of cells were lysed in buffer (25mM Tris-HCl pH=7.5, 500mM NaCl, 15mM imidazole pH=8.0) with 1mM PMSF and 1% DDM (Anatrace) via probe sonication. Lysed cells were solubilized for 1hr at 4°C and the centrifuged supernatant was loaded twice on a 3mL Ni-NTA column and washed with the same buffer. After washing with 10CVs lysis buffer +15mM Imidazole, pH=8.0, bound MolBC was eluted in 10CVs elution buffer +235mM Imidazole, pH=8.0 and concentrated using a 100kDa MWCO concentrator (Millipore). The protein was then injected over a Superdex 200 increase 10/300 GL (Cytiva) pre-equilibrated in buffer (25 mM Tris-HCl, pH 7.5, 300 mM NaCl, 0.02% DDM). SEC fractions were assessed by SDS-PAGE and assembled with MolA or stored at –80°C for other experiments.

### cryo-EM sample preparation and data collection

All cryo-EM grids were prepared in a Vitrobot Mark III set to 95-100% humidity and 4°C and loaded on glow-discharged Quantifoil holey carbon grids (R1.2/1.3μm Au, 300 mesh) with graphene coating (Atomface, formerly known as Beijing Graphene Institute Co., LTD). For the apo state, 4μL of MolBC at 0.5mg/mL was let absorb for 10sec and blot for 3sec with no applied blot force before plunge-frozen in liquid ethane pre-cooled in LN_2_. MolBC-A at 6mg/mL was similarly vitrified after complex assembly at a 1:1.7 MolBC:MolA ratio for 30min on ice, and excess MolA removal by SEC. All nucleotide-bound states employed the MolBC_E154Q_ assembled with MolA in presence of 2mM AMP-PNP/Mg^2+^ over SEC, 2mM ATP/Mg^2+^/Na_2_WO_4_ + 0.2mM Na_3_VO_4_ added before grid freezing, or 2mM ADP/Mg^2+^ added also prior to grid freezing. Na_3_VO_4_ was heated for 2min at 95°C to maintain its active form before mixing it with the complex and grid freezing. For the ATP state with Na_2_WO_4_, M9 grown MolA was used for complex assembly. To avoid particle crowding these samples were frozen at 0.4-0.6mg/mL and to increase ice thickness blotting was decreased to 2sec and blot force was set to –1. All cryo-EM datasets were collected remotely at the S2C2 Stanford-SLAC CryoEM Center in a Titan Krios (FEI) electron microscope set to 300kV, a nominal magnification of 105kx, and a 20eV slit width energy filter set in EPU software. Movies were recorded using a Gatan K3 direct electron detector with a binned pixel size of 0.86Å, a defocus range of –1.2 to –2.2μm, and a total dose of 50e^-^/Å^2^. Parameters varied slightly for ADP-bound MolBC-A that was collected at 1.086Å and a total dose of 60e^-^/Å^2^, and MolBC that had a defocus range of –0.5 to –2.5μm.

### cryo-EM data processing

All collected datasets were processed in CryoSPARC v4.7.1+. Detailed processing parameters for each dataset can be found in the **Supplementary Figures 4-8**, and data collection, refinement, and validation statistics can be found in the **Supplementary Table 1**. Imported movies were first patch motion corrected, and their contrast transfer function (CTF) was estimated. The CTF estimated statistics for each micrograph were then used to manually curate the exposures and exclude bad images from processing. Following micrograph denoising, initial 2D classes were generated on blob picked particles, Fourier cropped to ½ or ¼ of the original box size. Classes of different protein orientations were used to create an Ab-initio consensus volume prior to improving its poses and shifts by heterogeneous refinement to use as template for particle picking. Blob and template picked particles were then combined and further refined by rounds of heterogeneous refinement using as input the same consensus volume and 4-5 “dummy” volumes reconstructed by Ab-initio on the excluded 2D classes. Once the particle percentage assigned to the consensus volume reached approximately >80%, the particles were extracted without Fourier cropping and separated from any remaining junk through one last round of refinement with forced classification on. To improve map statistics, the particles were then non-uniform refined. C1 symmetry was applied in all the SBP-bound structures mimicking nucleotide-free or nucleotide-bound states, while C2 symmetry was applied only in the case of nucleotide-free MolBC. In cases of sub-optimal orientation diagnostics rebalancing of orientations was performed to improve non-uniform refinement results. Similarly, reference-based motion correction (RBMC) was applied in cases where the final map quality was improved. Final volumes and particle stacks were used downstream for model building or further refined to determine their conformational variability.

### cryo-EM model building and visualization

ModelAngelo, an automated model building tool, was used for atomic model reconstruction taking as input each 3D high-resolution cryo-EM map and a FASTA file including the protein sequence of each protein domain, as obtained from Uniprot^41^. The resulting structural models were inspected in ChimeraX-1.10 prior to import in Coot-0.9.8.92 (CCP4) for manual refinement^42,43^. Automated refinement was performed by iterative rounds of real-space refine in Phenix-2.0 with default settings and resolution specified^44^. Each round was inspected and further refined manually in Coot until no further improvements could be made and appropriate MolProbity scores were obtained. To measure the pore dimensions, each model was run through HOLE following deletion of the SBP and the NBDs and TM5/5α based alignment of the TMDs. Pore radius was plotted against the distance along the z-axis and the raw HOLE sphere files were used to visualize the results as a dot surface in PyMOL 3.1.4.1^45^.

### MD Simulations

Atomic models of the MolA substrate binding protein in the closed conformation with bound substrate (molybdate or tungstate) were constructed based on the high-resolution X-ray crystal structures of the molybdate-bound (PDB ID: 3PSH) and tungstate-bound (PDB ID: 3PSA)^35^. Additionally atomic models for MolA in the more open conformation (transporter-bound) with bound substrate were constructed based on the MolA-substrate free cryo-EM structure of the MolBC-A complex from this work. The substrates were placed in the binding pocket of the more open MolA conformation by superimposing the closed conformation with its bound substrate. The coordinating water molecule between K221 for tungstate-bound and Y114 for molybdate-bound were kept for both closed and more open MolA conformations. The MolA with bound substrate were solvated in 150mM NaCl using *Solution Builder* in CHARMM-GUI^46–48^. All residues were assigned their standard protonation state at pH=7. The total number of atoms in the model is about 91,000 atoms. The CHARMM36m force field for protein^49–52^, ions^53^, TIP3P water^54^, and parameters for tungstate and molybdate were based in combination on homology from CGenFF version 4.6^55,56^, Rmin/2 from^57^, and bonds and angles were assigned their average value from the X-ray crystal structures^35^. Then four instances of MD simulations were performed using OpenMM version 8.1.1 either with harmonic restraints on Cα atoms at 1kcal/mol/Å^2^ or no restraints^58^. All simulations were performed with periodic boundary conditions with long-range electrostatics treated by the particle mesh Ewald method^59^, a cutoff of 12Å, a switching function starting at 10Å, and all bonds involving hydrogen atoms held fixed. The simulations use a time step of 2fs. The temperature was held constant at 310K with the LangevinIntegator^60^ with a 1/ps friction coefficient. The pressure was held constant at 1atm with the MonteCarloBarostat^61,62^ with pressure changes attempted every 25 steps. Each simulation instance was performed as follows: The model was first energy minimized for 400 steps. Then equilibration was done for 100ps at NVT (constant number of particle N, volume V, and temperature T) conditions followed by 1.1ns at NPT (constant number of particle N, pressure P, and temperature T) conditions. The substrate was then pulled out of the binding pocket at a rate of 1Å/ns over the course of 15ns. The pulling was done by moving the restraint point of a radial harmonic restraint on the COM to COM (center of mass) distance between the backbone atoms and the substrate atoms beginning at about 5-6Å with a force constant of 1(kcal/mol/Å^2^)^63^.

### ATPase activity assays

The NADH-coupled ATPase assay was used to determine the ATPase activity of MolBC-A in the presence of MolA and substrate. The assay was set up in triplicate in a 384-well non-binding plate (Greiner, #781900). Each replicate consisted of 27uL of reaction mixture containing MolBC at 0.045μM final concentration pre-assembled with MolA at 0.090μM final concentration and grown in M9 media to remove any endogenous substrate levels. The complex was incubated for 30min on ice before molybdate or tungstate was introduced in a range of concentrations (0-6100μM; 2.5-fold serial dilution). The remaining mixture had 250μM NADH, 4mM phosphoenolpyruvate (PEP), 12-20U of pyruvate kinase (PK), and 18-28U of lactate dehydrogenase (LDH) added. A standard NADH curve was also prepared to calculate ATPase rates. Following plate incubation at 37°C for 5min, 2mM ATP was added and the decrease in NADH concentration was measured by absorbance at 340nm in a SPARK microplate reader (Tecan) in 30s intervals for 20min. Using linear regression analysis, the slopes for each condition were determined and subtracted from buffer background. Same analysis was used to calculate the rate of NADH disappearance from the standard curve and convert the changes in fold fluorescence (ΔRFU/s) to ATPase rates (nmol/mg/min).

### Particle orientation refinement using GMM

To explore the conformational landscape of MolA in MolBC-A in different states, we used e2gmm, a deep neural network (DNN) and Gaussian mixture model (GMM) based method for protein structural heterogeneity analysis^64^. Starting from the CryoSPARC refinement of each dataset, GMM representations were built for the volumes and global refinement were performed to improve particle orientation alignments^65^. Since MolA exhibited the highest level of flexibility in all maps, we then ran two rounds of focused refinement to improve density around MolA (round 1) and MolA Lobe II (round 2). The focused refinement results were then combined with the base global refinement to generate the final composite map for better structure interpretation. To characterize the type of motion MolA exhibited relative to the rest of the complex, motion trajectories of MolA were built from the orientation assignment of the focused refinements. The per-particle orientation difference between the global and MolA focused refinement were computed and mapped to a low dimensional space, and the dominant motion trajectory was visualized by reconstructing particles at different positions along the first eigenvector in the low dimensional space. Detailed parameters for each composite map can be found in the **Supplementary Table 2**.

### BLI Octet measurements and data analysis

To study the effect of substrate and nucleotide on SBP-transporter interactions, MolA was expressed in M9 media and purified to remove the His tag. Binding experiments were collected in triplicate in 25mM Tris-HCl, pH 7.5, 300 mM NaCl, 0.02% DDM at 25°C. MolBC was immobilized at 0.03mg/mL on a Ni-NTA Biosensor (Fisher Scientific, Sartorius). After establishing baseline for 30sec, each sensor was loaded for 300sec followed by 30sec dissociation to ensure complete and stable coating. To determine the MolA-MolBC affinity in absence of nucleotide and substrate, MolA (0-80nM; 2-fold serial dilution) was associated for 180sec and dissociation was monitored for 240sec. To determine the affinity of MolA for MolBC in the presence of ATP/Mg^2+^ or ADP/Mg^2+^, MolBC was immobilized in buffer containing 2mM final concentration of either nucleotide. Then, MolA (0-80nM; 2-fold serial dilution) was left to associate for 180sec followed by dissociation of 240sec in buffer containing either nucleotide at 2mM throughout all steps of the binding assay. For kinetic experiments with substrate, MolA at 10nM was associated with MolBC for 180sec to form a complex, before molybdate or tungstate (0-1mM; 2.5-fold serial dilution) was added only during dissociation and monitored for 240sec. Similarly, to test the combined effect of substrate and nucleotide on MolA-MolBC interactions, MolBC was immobilized in buffer containing 2mM final ATP/Mg^2+^. MolA at 10nM was then associated with MolBC in presence of nucleotide for 180sec and molybdate or tungstate (0-1mM; 1.5-fold serial dilution) was added only during dissociation for 240sec in buffer containing ATP/Mg^2+^. For each binding experiment, non-specific binding tests of each analyte were performed to establish background. Data were collected on the Octet K2 two-channel instrument (Pall Life Sciences). All binding data were analyzed in the Data Analysis HT 11.0.0.50 (Sartorius) software. Preprocessed data were background corrected using appropriate reference samples and the y-axis was aligned to the baseline average. Inter-step correction was aligned to the association step to account for baseline drifts that were minimal. Kinetic parameters in absence of substrate were determined by analyzing association and dissociation steps and data were fit using a 1:1 binding model. Kinetic parameters in presence of substrate (molybdate or tungstate) were determined by performing a dissociation only fit.

## Data Availability

The cryo-EM maps of the wild-type MolBC, MolBC-A, MolBC-A(E154Q) bound to ATP, AMP-PNP, and ADP have been deposited in the Electron Microscopy Data Bank under accession numbers: MolBC^WT^ (EMB-78939), MolBC-A^WT^ (EMB-79077), MolBC-A^E154Q^ ATP/Mg^2+^ (EMB-78941), MolBC-A^E154Q^ AMP-PNP (EMB-79076), and MolBC-A^E154Q^-ADP (EMB-79078). Atomic coordinates for the atomic models have been deposited in the Protein Data Bank [http://www.rcsb.org] under accession numbers PDB ID: MolBC^WT^ (PDB-38MO), MolBC-A (PDB-38RC), MolBC-A^E154Q^ATP/Mg^2+^ (PDB-38MT), MolBC-A^E154Q^ AMP-PNP (PDB-38RB), and MolBC-A^E154Q^-ADP (PDB-38RD). Composite maps from the GMM-based focused refinement of MolBC-A(E154Q) bound to ATP, AMP-PNP, and ADP have also been deposited in the Electron Microscopy Data Bank under accession numbers: MolBC-A^E154Q^ ATP/Mg^2+^ (EMB-79108), MolBC-A^E154Q^ AMP-PNP (EMB-79113), and MolBC-A^E154Q^-ADP (EMB-79121). Atomic coordinates for the atomic models out of heterogeneity analysis have been deposited in the Protein Data Bank [http://www.rcsb.org] under accession numbers PDB ID: MolBC-A^E154Q^ATP/Mg^2+^ (PDB-38SL), MolBC-A^E154Q^ AMP-PNP (PDB-38SO), and MolBC-A^E154Q^-ADP (PDB-38SV). All data are available in the main text, the Supplementary Information or submitted to Figshare.

## Acknowledgments

The authors acknowledge the University of Chicago Research Computing Center for computational resources provided by the Beagle 3 high-performance computing cluster, supported by the National Institutes of Health (NIH) under grant 1S10OD028655-01. The authors acknowledge the Keck Biophysics Facility, a shared resource of the Robert H. Lurie Comprehensive Cancer Center of Northwestern University, supported in part by the NIH NCI Cancer Center Support Grant P30 CA060553. Molecular graphics and analyses were performed with UCSF ChimeraX, developed by the Resource for Biocomputing, Visualization, and Informatics at the University of California, San Francisco, with support from NIH R01-GM129325 and the Office of Cyber Infrastructure and Computational Biology, National Institute of Allergy and Infectious Diseases (NIAID). Finally, the authors thank, Evan Buechel, Valentina Dimitrova, Anna Gilstrap, Lydia Kenney and Chloe Jackson for manuscript comments and proofreading.

## Funding

H.W.P acknowledges support from the National Institutes of Health (R35GM158705 and R01GM140584); B.R. acknowledges support from the National Institutes of Health (R35-GM152124); M.C. acknowledges support from the National Institutes of Health (R01GM150905).

## Author information

Department of Molecular Biosciences, Northwestern University, Evanston, IL 60208, USA.

Alexandra Karagiaridi and Heather W. Pinkett

Department of Chemistry, University of Chicago, Chicago, IL 60637, USA.

Ramon Mendoza Uriarte

Division of CryoEM and Bioimaging, SSRL, SLAC National Accelerator Laboratory, Stanford University, Menlo Park, CA 94025, USA.

Muyuan Chen

Department of Biochemistry and Molecular Biology, University of Chicago, Chicago, IL 60637, USA.

Benoît Roux

## Author Contribution

A.K performed MolA and MolBC purifications, cryo-EM data collection and single-particle cryo-EM analyses as well as BLI and ATPase experiments. M.C carried out the particle orientation refinement using GMM. R.U and B.R. carried out the MD simulations. A.K. and H.P. wrote the manuscript. H.P. conceived and supervised the work.

## Ethics declaration

The authors declare no competing interests.

## Notes

### Competing Interest Statement

The authors have declared no competing interest.

