## Supplemental for "Nucleotide State Regulates Asymmetric Substrate-Binding Protein Docking in a Type II ABC Importer"

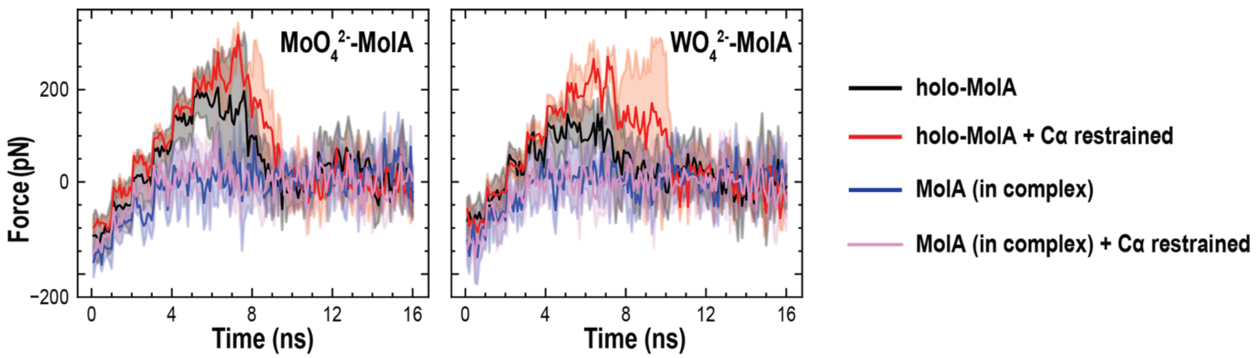


**Supplementary Figure 1: Remodeling of MolA into a more open conformation requires less force to unbind substrate from the binding pocket.** Plots of the force required to unbind molybdate (left) or tungstate (right) from MolA either in complex or isolated (holo-MolA) with (pink, red) and without (blue, black) restraints, over the 15ns simulation. Each trace is the average of four independent simulations (n=4), with standard deviation represented by the gray shaded area.

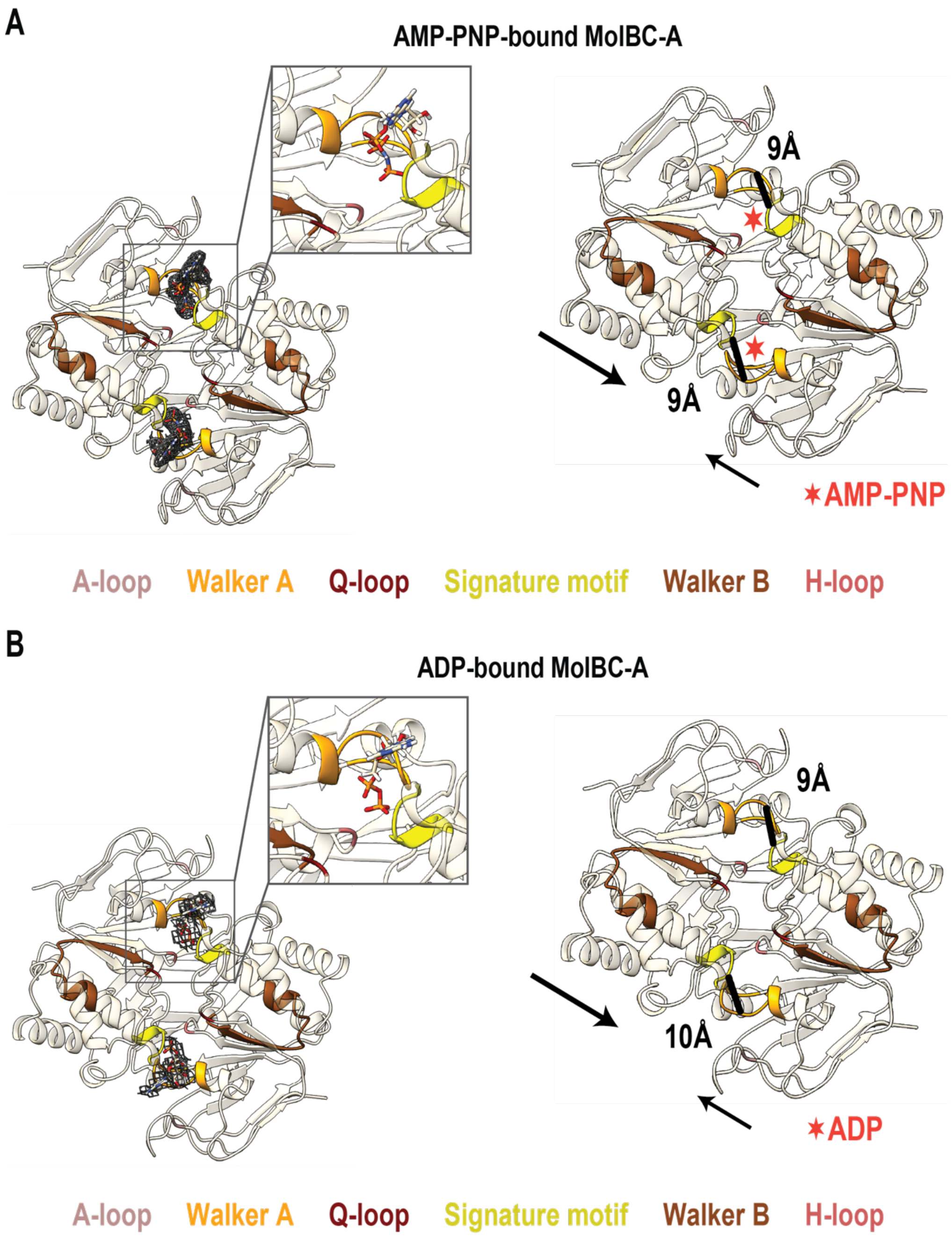


**Supplementary Figure 2: AMP-PNP and ADP each induces asymmetric closure of the NBDs comparable to ATP. A, Left,** Enlarged top view of the AMP-PNP binding sites in AMP-PNP-bound MolBC-A. AMP-PNP shown as sticks, with corresponding density shown around each ligand. Conserved nucleotide-binding motifs are colored according to the key. **Right,** Cytoplasmic (top) views of the dimerized MolC NBDs (wheat) in AMP-PNP-bound MolBC-A, showing AMP-PNP-induced NBD closure. Cα-Cα distances between S130 in the signature motif of one MolC subunit and C42 in the Walker A motif of the opposing subunit are indicated for both NBDs. Arrows indicate the unequal movements of the two MolC subunits upon AMP-PNP binding. Red asterisks mark the positions of AMP-PNP, which are omitted for clarity. **B,** Same views as in **A** but for ADP-bound MolBC-A.

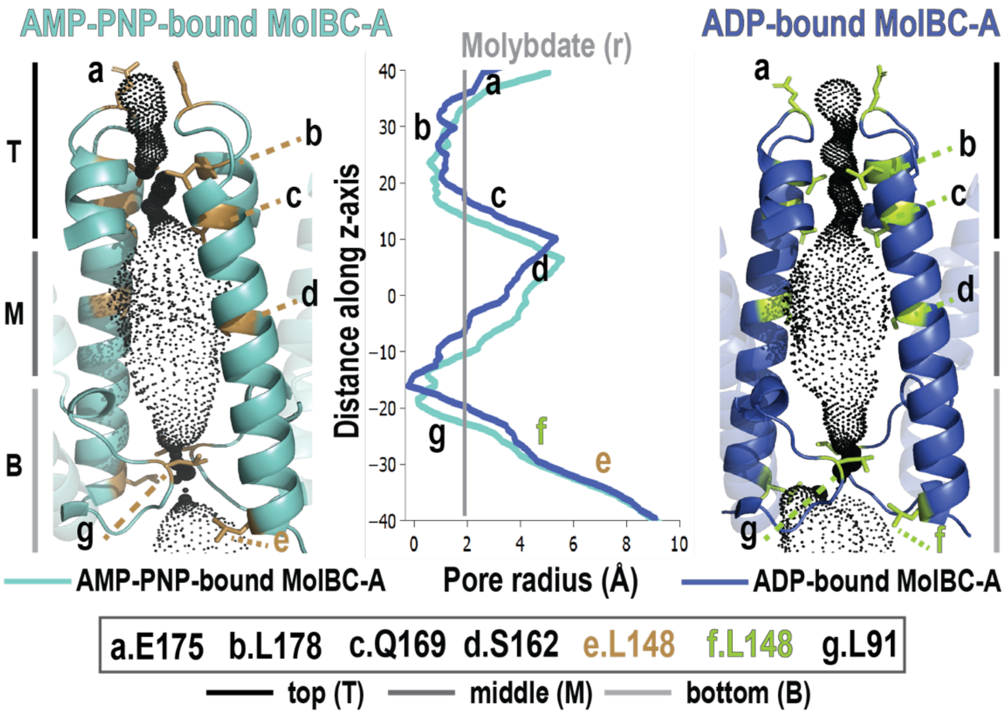


**Supplementary Figure 3: AMP-PNP and ADP binding each remodels the transport pathway similarly to ATP.** Comparison of the translocation pathway of AMP-PNP-bound MolBC-A (left; light blue) and ADP-bound MolBC-A (right; dark blue). TM5 and TM5α, which line the central pathway, are shown in cartoons, with the pore dimensions calculated using HOLE represented by black points. The central plot shows pore radius along the membrane-normal z-axis from the periplasmic (top) to the cytoplasmic side (bottom). E175 marks the periplasmic entrance, L178 the deeper TM5α constriction, Q169, and S162 the midpoint of the pathway, and L148 and L91 cytoplasmic gates I and II, respectively.

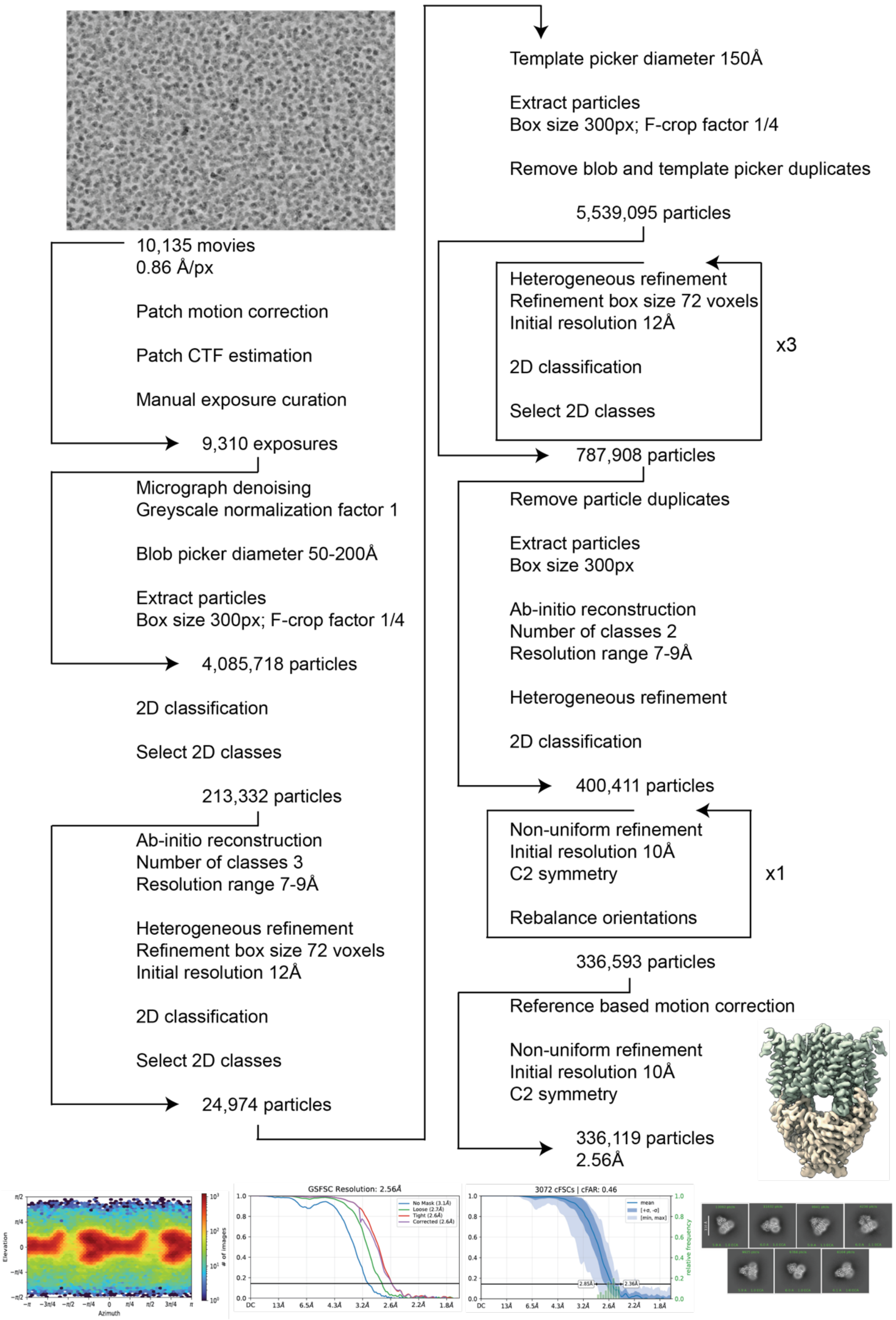


**Supplementary Figure 4: cryo-EM processing flowchart for recombinantly expressed nucleotide-free (apo) MolBC from *H. influenzae* in 0.02% DDM.**

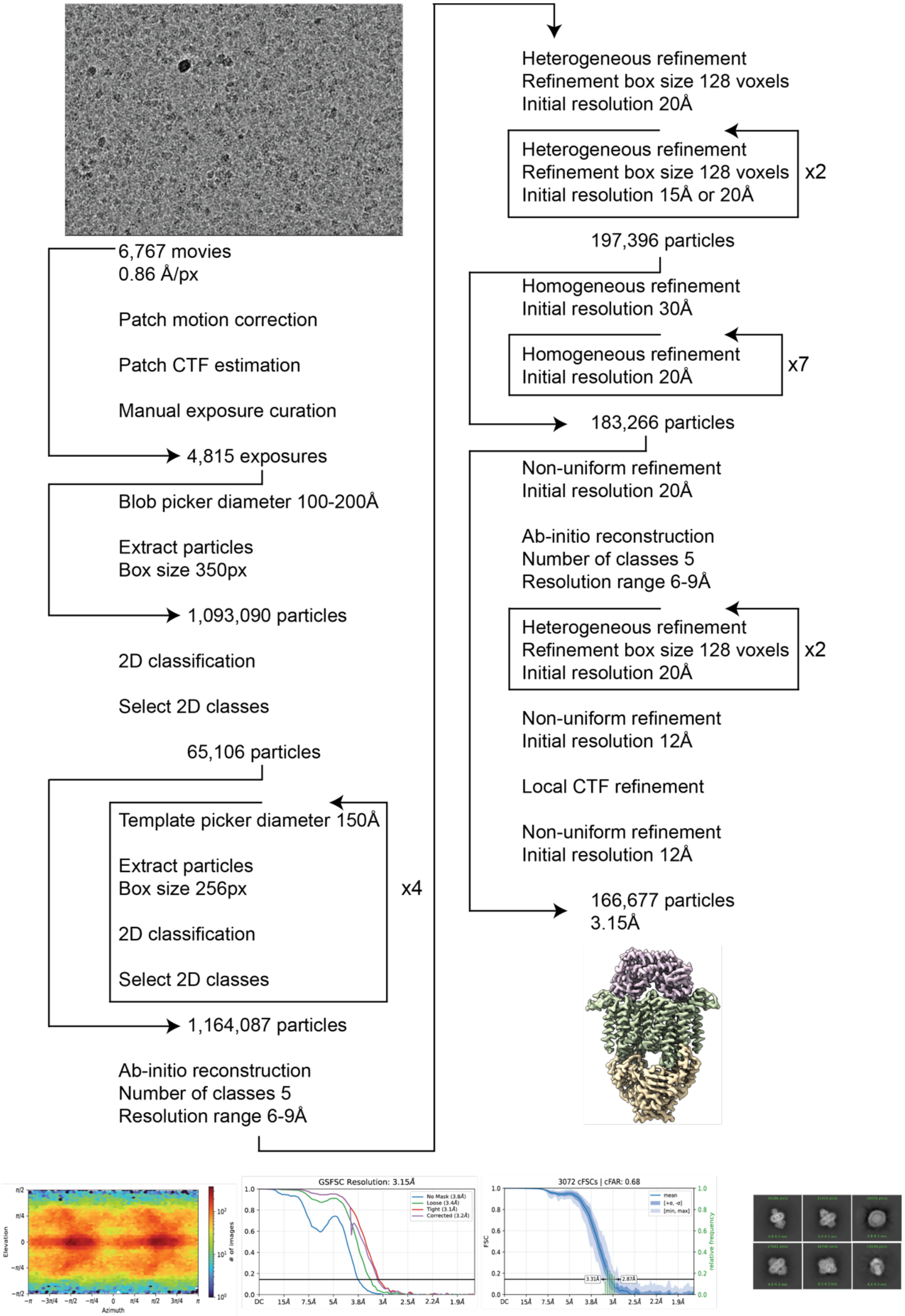


**Supplementary Figure 5: cryo-EM processing flowchart for recombinantly expressed nucleotide-free (apo) MolBC-A from *H. influenzae* in 0.02% DDM.**

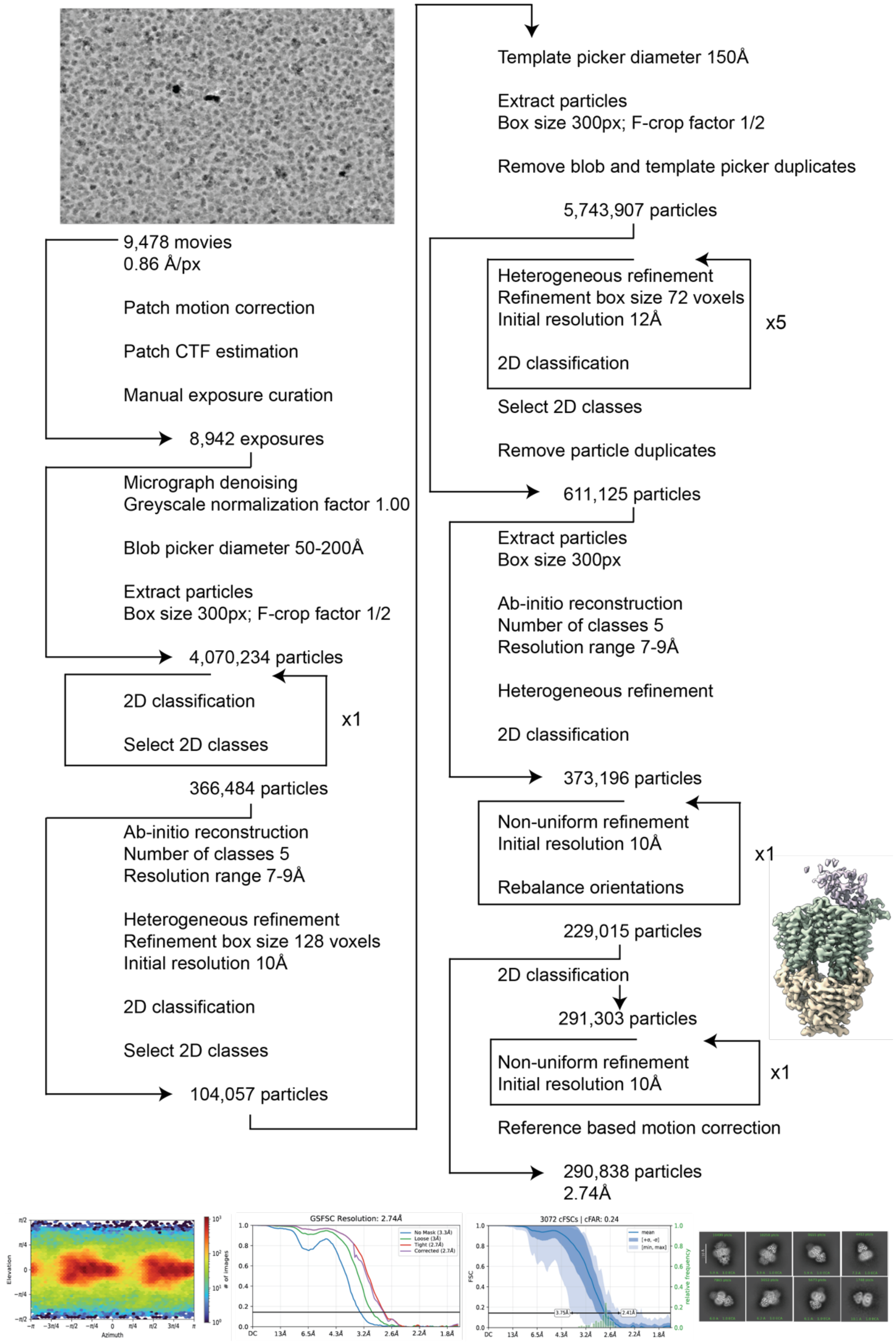


**Supplementary Figure 6: cryo-EM processing flowchart for recombinantly expressed ATP-bound MolBC-A from *H. influenzae* in 0.02% DDM.**

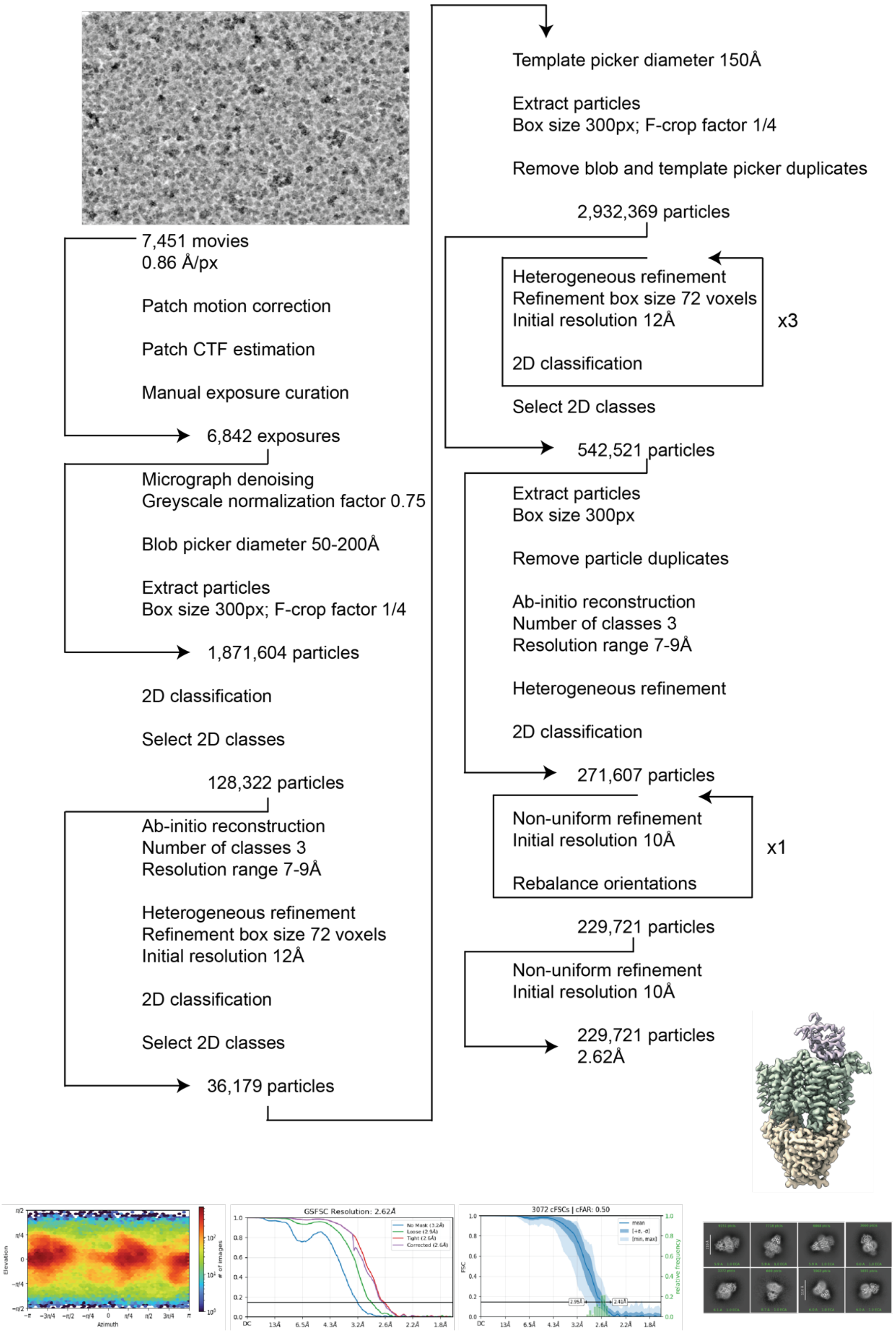


**Supplementary Figure 7: cryo-EM processing flowchart for recombinantly expressed AMP-PNP-bound MolBC-A from *H. influenzae* in 0.02% DDM.**

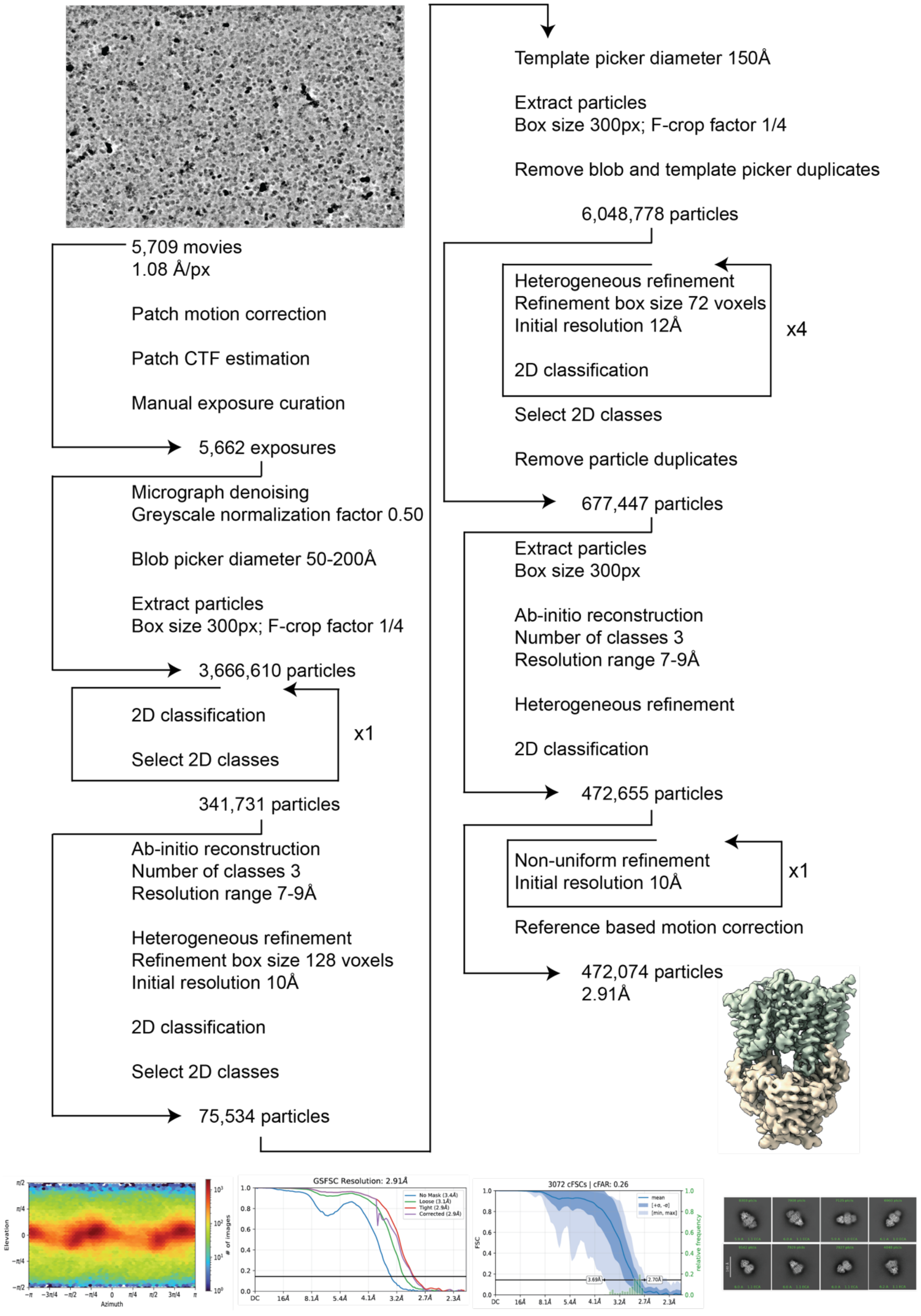


**Supplementary Figure 8: cryo-EM processing flowchart for recombinantly expressed ADP-bound MolBC-A from *H. influenzae* in 0.02% DDM.**

**Supplementary Table 1: cryo-EM data collection, refinement, and validation statistics.**

| | | | | | |
|---|---|---|---|---|---|
| **Data Collection** | | | | | |
| Magnification | | | 105,000 | | |
| Voltage (kV) | | | 300 | | |
| Energy filter width (eV) | | | 20 | | |
| Maps | MolBC | MolBC-A | MolBC-A + AMP-PNP | MolBC-A + ATP | MolBC-A + ADP |
| Defocus range (μm) | -0.5 to -2.5 | -1.3 to -2.3 | -1.2 to -2.2 | -1.2 to -2.2 | -1.2 to -2.2 |
| Pixel size (Å) | 0.86 | 0.86 | 0.86 | 0.86 | 1.08 |
| Total dose ($e^-$/Å$^2$) | 50 | 50 | 50 | 50 | 60 |
| Number of micrographs | 10,135 | 6,767 | 7,451 | 9,478 | 5,709 |
| Number of initial particles | 4,085,718 | 1,093,090 | 1,871,604 | 4,070,234 | 3,666,610 |
| **Refinement** | | | | | |
| Number of final particles | 336,119 | 166,677 | 229,721 | 290,838 | 472,074 |
| FSC threshold | 0.5 | 0.5 | 0.5 | 0.5 | 0.5 |
| GSFSC resolution (Å) | 2.56 | 3.15 | 2.62 | 2.74 | 2.91 |
| Symmetry | C2 | C1 | C1 | C1 | C1 |
| *B* factor (Å$^2$) | -105.7 | -105.1 | -86.5 | -106.7 | -118.7 |
| **Model composition** | | | | | |
| Chains | 4 | 5 | 5 | 5 | 4 |
| Non-hydrogen atoms | 8,803 | 11,425 | 10,073 | 9,329 | 8,347 |
| Protein residues | 1,131 | 1,463 | 1,284 | 1,190 | 1,067 |
| Ligand | 0 | 0 | 2 | 4 | 2 |
| R.M.S deviations | | | | | |
| Bond lengths (Å) | 0.003 | 0.003 | 0.003 | 0.004 | 0.003 |
| Bong angles (°) | 0.573 | 0.538 | 0.553 | 0.697 | 0.672 |
| **Validation** | | | | | |
| MolProbity score | 1.14 | 1.13 | 1.05 | 1.72 | 1.66 |
| Clash score | 2.39 | 1.59 | 2.39 | 5.69 | 6.30 |
| Poor rotamers (%) | 0.92 | 0.40 | 0.45 | 0.97 | 0.98 |
| Ramachandran plot | | | | | |
| Favored (%) | 97.41 | 96.55 | 97.87 | 93.92 | 95.46 |
| Allowed (%) | 2.41 | 3.45 | 1.97 | 5.48 | 3.97 |
| Disallowed (%) | 0.18 | 0.00 | 0.16 | 0.60 | 0.57 |

**Supplementary Table 2: cryo-EM data collection, refinement, and validation statistics for each focused refined map in EMAN2.**

| | | | |
|---|---|---|---|
| **Model composition** | | | |
| Maps | MolBC-A + AMP-PNP | MolBC-A + ATP | MolBC-A + ADP |
| Chains | 5 | 5 | 4 |
| Non-hydrogen atoms | 10,547 | 10,607 | 8,407 |
| Protein residues | 1,343 | 1,349 | 1,073 |
| Ligand | 2 | 4 | 2 |
| R.M.S deviations | | | |
| Bond lengths (Å) | 0.003 | 0.004 | 0.004 |
| Bong angles (°) | 0.559 | 0.637 | 0.695 |
| **Validation** | | | |
| MolProbity score | 1.42 | 1.66 | 1.88 |
| Clash score | 2.79 | 4.96 | 5.49 |
| Poor rotamers (%) | 1.73 | 1.04 | 2.28 |
| Ramachandran plot | | | |
| Favored (%) | 97.05 | 94.22 | 95.58 |
| Allowed (%) | 2.72 | 5.55 | 4.05 |
| Disallowed (%) | 0.23 | 0.23 | 0.38 |
